# The nuclear effector ArPEC25 from the necrotrophic fungus *Ascochyta rabiei* targets the chickpea transcription factor CaβLIM1a and negatively modulates lignin biosynthesis for host susceptibility

**DOI:** 10.1101/2021.09.02.458738

**Authors:** Shreenivas Kumar Singh, Sandhya Verma, Kunal Singh, Ankita Shree, Ritu Singh, Vikas Srivastava, Kamal Kumar, Ashutosh Pandey, Praveen Kumar Verma

## Abstract

Fungal pathogens deploy a barrage of secreted effectors to subvert host immunity, often by evading, disrupting, or altering key components of transcription, defense signaling, and metabolic pathways. However, the underlying mechanisms of effectors and their host targets are largely unexplored in necrotrophic fungal pathogens. Here, we describe the effector protein ArPEC25, which is secreted by the necrotroph *Ascochyta rabiei*, the causal agent of Ascochyta blight disease in chickpea (*Cicer arietinum*), and is indispensable for virulence. After entering host cells, ArPEC25 localizes to the nucleus and targets the host LIM transcription factor CaβLIM1a. CaβLIM1a is a transcriptional regulator of *CaPAL1*, which encodes phenylalanine ammonia lyase, the regulatory, gatekeeping enzyme of the phenylpropanoid pathway. ArPEC25 inhibits the transactivation of CaβLIM1a by interfering with its DNA binding ability. This results in negative regulation of the phenylpropanoid pathway and decreased levels of intermediates of lignin biosynthesis, thereby suppressing lignin production. Our findings illustrate the role of fungal effectors in enhancing virulence by targeting a key defense pathway that leads to the biosynthesis of various secondary metabolites and antifungal compounds. This study provides a template for the study of less explored necrotrophic effectors and their host target functions.

**One-sentence summary:** The *Ascochyta rabiei* effector ArPEC25 enters the host nucleus and targets the transcription factor CaβLIM1a to manipulate phenylpropanoid pathway for negative modulation of chickpea lignin biosynthesis.

## Introduction

Agricultural crops are continually exposed to biotic factors (pathogens) that can cause severe economic losses. Plant fungal pathogens have broadly evolved into two groups, as defined by their infection cycles: biotrophs (with a predominantly biotrophic phase) and necrotrophs (with a predominantly necrotrophic phase) (Seybold et al., 2020). During their pathogenesis and proliferation, biotrophs maintain a tightly regulated interaction with their hosts that keeps them alive, whereas necrotrophic fungal pathogens promote necrosis and the death of their hosts to feed on the released nutrients (Ökmen and Doehlemann, 2014; Mengiste, 2012). Nevertheless, results gathered from the necrotrophs *Botrytis cinerea* and *Sclerotinia sclerotiorum* have suggested that necrotrophs have a short biotrophic phase during the early stages of infection (Williams et al., 2011; Shlezinger et al., 2011; Seifbarghi et al., 2017; Rajarammohan, 2021). Instead of arbitrarily killing their hosts, necrotrophs, like biotrophs, elegantly manipulate crucial biological processes in their hosts to delay cell and tissue necrosis. Only at later stages does the infection enter the necrotrophic phase, which results in the onset of cell death to nourish the pathogen (Veloso et al., 2018).

Invading plant pathogenic fungi and oomycetes secrete an arsenal of specialized molecules called effectors that facilitate successful infection. The current view of plant-pathogen interactions suggests that the effectors secreted by biotrophic fungi and oomycetes circumvent host immunity by suppressing host cell death. Less is known about the effectors secreted by necrotrophic pathogens, with the exception of cell wall–degrading enzymes that promote host susceptibility and cell death (Liu et al., 2019a). A compatible interaction between a host-specific receptor and its cognate effector induces the onset of effector-triggered immunity (ETI) in biotrophic interactions. By contrast, the direct or indirect interaction of host-specific proteins with necrotrophic effectors (NEs) often triggers cell death, which culminates in host susceptibility (effector-triggered susceptibility; ETS) (Sung et al., 2021; Oliver and Solomon Peter S, 2010). The victorin–LOCUS ORCHESTRATING VICTORIN EFFECTS 1 (LOV1) interaction that occurs during infection of Arabidopsis (*Arabidopsis thaliana*) by the necrotroph *Cochliobolus victoriae* is one of the few known examples of effector-target interaction leading to susceptibility (Winterberg et al., 2014). Similarly, the secreted effector SnTox3 from *Parastagonospora nodorum* facilitates disease progression in wheat (*Triticum aestivum*) upon interaction with its receptor Pathogenesis-related 1 (TaPR1) (Sung et al., 2021; Breen et al., 2016). Two other examples of interactions between NEs and their cognate host targets studied in the same pathosystem are SnToxA-Tsn1 (Liu et al., 2006) and SnTox1-Snn1 (Liu et al., 2004). Likewise, only three interactions between NEs and their host targets have been characterized for their cell death phenotype upon pathogen attack in the *Pyrenophora tritici-repentis*-wheat pathosystem: PtrToxA-Tsn1, PtrToxB-Tsc2, and PtrToxC-Tsc1 (Corsi et al., 2020).

The current consensus about these pathosystems is that the pathogen secretes effector proteins to circumvent the host innate immunity pathway. Although the effectors secreted by pathogenic microbes are extremely diverse, they sometimes carry a conserved N-terminal amino acid sequence that plays a role in effector secretion and translocation. The few examples of characterized motifs include the RxLR, LFLAK-HVLVxxP, Crinkler (CRN), Y/F/WxC, CFEM, LysM, DELD, EAR, and RGD motifs (Snelders et al., 2020; Boddey et al., 2016). The RxLR motifs are well characterized for their role in the virulence of various oomycete phytopathogens (Liu et al., 2019b). Additionally, the recent availability of genome sequences for various phytopathogenic fungi has revealed the presence of a fifth conserved amino acid residue in the RxLR motif that suggested its high similarity to the Plasmodium Export Element (PEXEL). PEXEL motifs from the malaria parasite (*Plasmodium* spp.) share the conserved sequence RxLxE/D/Q (with x being any amino acid), which is positioned close to the N-terminal secretory signal sequence (Luisa Hiller et al., 2004). This conserved sequence is often cleaved by the endoplasmic reticulum (ER) resident protease Plasmepsin V (PMV) (Boddey et al., 2016, 2010, 2009). This cleavage step is crucial for effector secretion from the parasite into host erythrocytes. The in silico comparative genome and secretome analyses of many phytopathogens suggested that fungal genomes encode more PEXEL-containing effectors than *Plasmodium* spp. (Choi et al., 2010).

A preformed physical barrier such as a thickened plant cell wall constitutes the first line of protection against most pathogens. To invade and colonize the host, attacking necrotrophs must overcome these physical barriers by penetrating directly through a natural opening or indirectly by using a penetration peg and by secreting cell wall–degrading enzymes (CWDEs) (Majd, 2007; Łaźniewska et al., 2009). Plants respond to necrotrophic attack by inducing the reinforcement of the cell wall through lignification. The phenylpropanoid biosynthetic pathway produces monolignols (G-, H-, and S-lignin), which are the building blocks of polymerized lignin. Genes participating in the phenylpropanoid pathway are strongly transcriptionally induced upon pathogen invasion and belong to multigene families, with phenylalanine ammonia lyase acting as a gateway enzyme (Bhuiyan et al., 2009; Zhang and Liu, 2015). Two-dimensional proteomic studies in the rapeseed (*Brassica napus*)-*Alternaria brassicae* pathosystem showed that cinnamyl alcohol dehydrogenase (CAD), which catalyzes the final step in the phenylpropanoid pathway specific to lignin formation, accumulates 48 h after pathogen infection (Sharma et al., 2007). The phytohormone salicylic acid (SA) provides resistance to a range of pathogens and is itself a product of the phenylpropanoid pathway, but its levels have been shown to be manipulated by secreted effectors from bacteria, oomycetes, and fungi (Shine et al., 2016; Yuan et al., 2019a; Liu et al., 2014, Bauters et al., 2021). Importantly, host susceptibility resulting from NEs specifically targeting the monolignol biogenesis branch of the phenylpropanoid pathway has not been reported yet. However, a few studies showed that the *Botrytis cinerea* elicitor BcGs1 and the *Parastagnospora nodorum* NE SnTox3 caused upregulation of the phenylpropanoid pathway and increased lignin deposition (Winterberg et al., 2014; Yang et al., 2018). The Tin2 effector protein of *U. maydis* indirectly modulates the phenylpropanoid pathway by rewiring metabolite flow into the anthocyanin pathway, which results in suppression of lignin accumulation (Brefort et al., 2014). Additionally, the Sta1 effector from the same pathogen suppresses the phenylpropanoid pathway, indicating modulation of lignin content (Tanaka et al., 2020). However, the molecular mechanism behind the modulation of lignin biogenesis by these NEs has not been completely deciphered.

The necrotrophic fungus *Ascochyta rabiei* causes Ascochyta blight (AB) disease and is a major constraint to chickpea (*Cicer arietinum*) production worldwide. Fungal conidia form germ tube– like structures that subsequently develop into an appressorium to penetrate host tissues (Fondevilla et al., 2015). Fungal hyphae grow sub-epidermally and produce necrotic lesions on chickpea leaves (Fondevilla et al., 2015). Several independent studies have examined the devastating effects of the pathogen on chickpea production (Viotti et al., 2012; Kaiser et al., 2000; Galdames and Mera, 2003). While breeding of genetically resistant chickpea cultivars has been attempted using quantitative trait loci (QTLs) for resistance to AB disease (Kumar et al., 2018), the fungal effectors involved in AB diseases have remained largely unexplored. Analyses of the *A. rabiei* genome, transcriptome, and secretome have revealed a variety of *A. rabiei* effectors with possible roles in pathogen virulence (Singh et al., 2012a; Fondevilla et al., 2015; Verma et al., 2016; Maurya et al., 2020; Mohd Shah et al., 2020). Earlier studies on *A. rabiei* pathogenesis suggested a role for the fungal toxins solanapyrones A, B, and C as well as cytochalasin D in virulence (Alam et al., 1989; Hamid and Strange, 2000). However, the deletion of the solanopyrone biosynthesis gene cluster demonstrated that these phytotoxins are not required for pathogenicity (Kim et al., 2015). Additionally, a recent study also proposed the involvement of endocytosis in *A. rabiei* virulence and a role for the membrane curvature sensing protein ArF-BAR in effector secretion (Sinha et al., 2021).

One of the most important mechanisms to suppress host defense employed by pathogens entails the targeting of fungal effectors to the host nucleus to reprogram the host transcriptional network. Several fungal effectors such as See1 (*Ustilago maydis*), PstGSRE1 (*Puccinia striiformis* f.sp. *tritici*), and CgEP1 (*Colletotrichum graminicola*) translocate to and function inside the host cell nucleus (Redkar et al., 2015; Qi et al., 2019; Vargas et al., 2016; Kim et al., 2020). Moreover, how nucleus-targeted effectors lead to host susceptibility is also poorly understood in fungi, especially in the case of necrotrophs. In this study, we report the identification and characterization of the PEXEL motif–containing nuclear effector ArPEC25 from *A. rabiei*, which is indispensable for virulence. Notably, ArPEC25 interferes with the activity of its host transcription factor target, CaβLIM1a, leading to susceptibility in chickpea. These findings pave the way to a better understanding of the mechanisms of effector action and provide new insights into necrotrophic fungal virulence.

## Results

### *ArPEC25* of *A. rabiei* is a potential effector candidate

Our earlier work on genome sequencing of the necrotrophic fungal pathogen *A. rabiei* (ArD2) revealed the presence of several putative secretory proteins (Verma et al., 2016). Motif search analysis showed that [Y/F/W]xC is the most frequent among all characterized motifs. The PEXEL motif initially identified in the malaria parasite *Plasmodium* spp. was also highly abundant, being present in 122 secretory proteins, in contrast to the RxLR motif (present in 38 proteins) commonly found among effectors of oomycete pathogens (Supplemental Table S1). In silico predictions for effector localization determined that 17 out of 122 candidate PEXEL motif–containing effectors harbor either a monopartite or a bipartite nuclear localization signal (NLS), suggesting nuclear localization in the host nucleus (Verma et al., 2016). We then analyzed transcript levels for the encoding fungal genes using data from our earlier differential transcript expression study using a suppression subtractive hybridization (SSH) library of *A. rabiei* upon oxidative stress: One gene, encoding the putative secretory protein ST47_g1734, was highly expressed. This protein, which possessed a typical PEXEL motif, was designated ArPEC25 (PEXEL-like Effector Candidate 25) and selected for further characterization (Singh et al., 2012b).

*ArPEC25* encodes a small cysteine-rich protein of 134 amino acids (aa). The SignalP 4.1 server predicted a cleavage domain of 28 aa at the N terminus of ArPEC25 (Figure 1A). The ArPEC25 protein lacked any known functional or structural domains other than the PEXEL motif (RTLND), which was located 11 aa downstream of the signal peptide (SP) cleavage site. The PEXEL motif was followed by an arginine-rich patch (RKRRRRR) (Figure 1A). Using ArPEC25 as a query, we identified ArPEC25 homologs in 29 fungal species representing diverse groups such as saprophytes, symbionts, animal pathogens, and plant pathogens with biotrophic, hemi-biotrophic, and necrotrophic lifestyles. Multiple alignments showed that the PEXEL motif and arginine-rich region are highly conserved in various fungal species (Supplemental Figure S3). These homologs also have two highly conserved cysteine residues that are characteristic features of many known effector proteins. A stretch of 10 residues (RECPVPKPGG) at the C terminus was also highly conserved.

**Figure 1.**
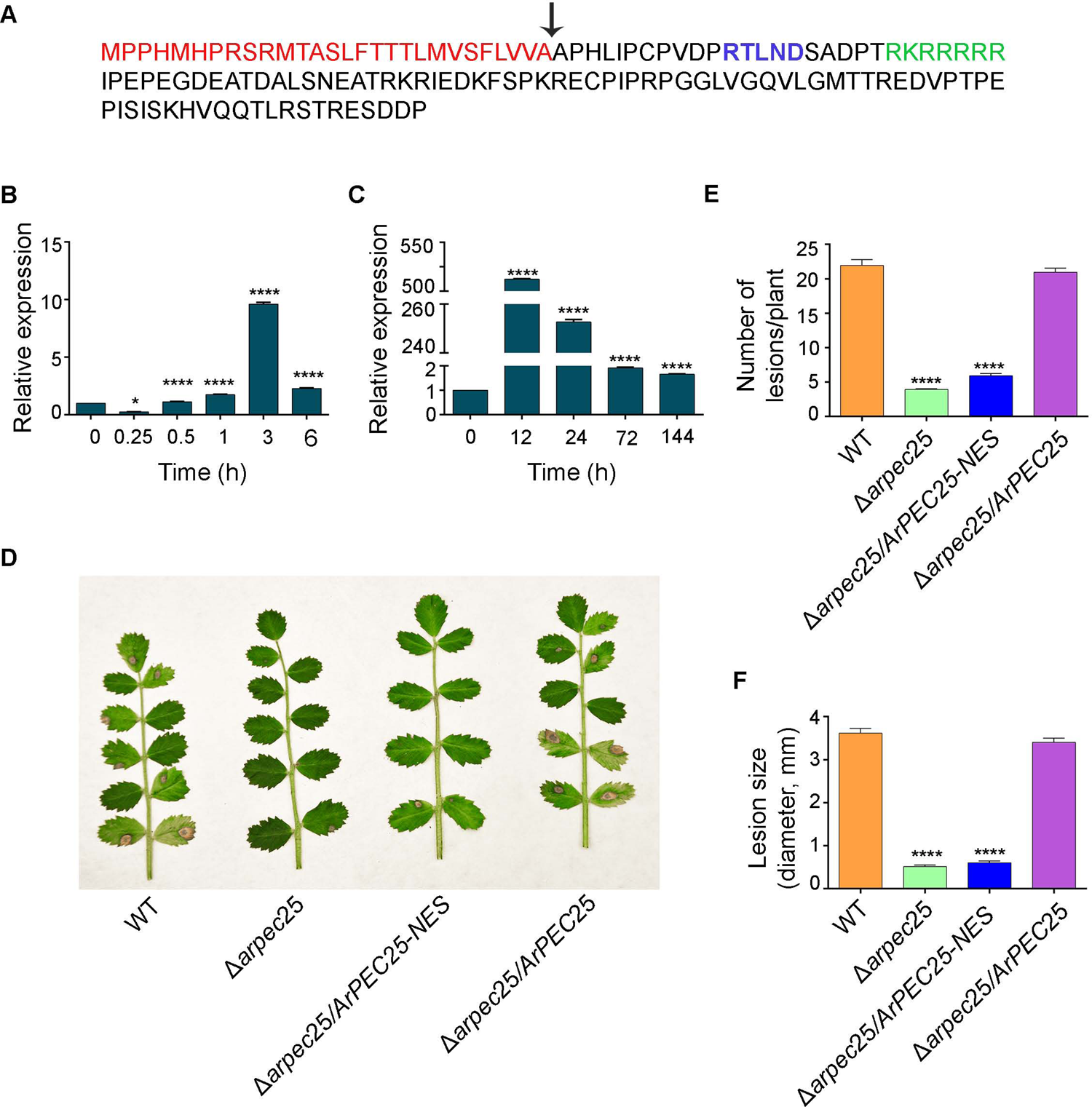
*ArPEC*25 is important for virulence of *A. rabiei*. **(A)** Schematic representation of amino acid sequence of ArPEC25 from wild type, *A. rabiei*. Signal peptide (SP) and its cleavage site is represented by red color and black arrow, respectively. Conserved PEXEL motif is shown in bold blue color and nuclear localization sequence is highlighted with green color. **(B)** Menadione induced *ArPEC25* expression level in wild type *A. rabiei*. mRNA was extracted from 250 µM menadione treated fungal mycelia at different time point for gene expression analysis. Ethanol treated tissues were taken as control and elongation factor *ArEF1α* (ST47_g4052) used for normalization. Data represents mean ± SD value of three independent experiments with three technical replicates each (n=3). Statistically significant differences were calculated by unpaired two tailed t-test; **** P≤0.0001 and * P ≤ 0.0160. **(C)** Expression analysis of *ArPEC*25 in wild type *A. rabiei* infected chickpea. mRNA was extracted from aerial tissues of infected chickpea at different time points for gene expression study. Transcript abundance was normalized with *ArEF1α*. Error bar represents ± SD from three independent biologicals replicates (n=3). Statistically significant differences were calculated by unpaired two tailed t-test; **** P ≤ 0.0001. **(D)** Image of chickpea Pusa362 at 7 days post inoculation (dpi) with wild type *A*. *rabiei* and mutant transformants. Image shows the necrotic lesions on leaves of chickpea plants infected with indicated transformants; WT, *Δarpec*25, *Δarpec*25/*ArPEC25*-*NES* and *Δarpec25*/*ArPEC25*, respectively. **(E)** and **(F)** Statistical analysis of lesion numbers and diameter on chickpea leaves infected with transformants described in **(D)**. The graph represents **(E)** mean number of lesions and **(F)** mean diameter of lesions from 50 plants. Bioassay was performed in three independent experiments with three technical replicates each. For **(E)** and **(F)**, the error bar represents ± SD of three independent experiments. Statistically significant differences were calculated by unpaired two tailed t-test; **** P≤0.0001.

### *A. rabiei*–secreted ArPEC25 accumulates during infection and is necessary for virulence

To determine the role of ArPEC25 during fungal infection, we generated gene deletion mutants (*Δarpec25*) via homologous recombination in the wild-type *A. rabiei* strain (Supplemental Figure S1A). Vegetative growth and conidiation of *Δarpec25* were comparable to those of the wild type (Supplemental Figure S1E). We confirmed that the *Δarpec*25 deletion strain carries only the single integration that disrupts *ArPEC25*, as determined by Southern blot hybridization and real-time PCR (Supplemental Figure S1, B and D). We performed bioassays on the chickpea cultivar ‘Pusa362’ with wild-type *A. rabiei* and the *Δarpec25* mutant, which exhibited markedly, reduced virulence compared to the wild type (WT) (Figure 1D and Supplemental Figure S1C). We also generated the complementation strain *Δarpec25*/*ArPEC25* to validate the association between loss of virulence and the loss of ArPEC25: Importantly, the wild-type copy of *ArPEC25* rescued virulence of the *Δarpec25* mutant. The complemented strain, *Δarpec25*/*ArPEC25*, was fully virulent and produced the characteristic blight disease in chickpea plants at rates similar to those of the WT (Figure1D and Supplemental Figure S1C). Bioinformatics analysis had suggested the host cell nucleus as the probable location of ArPEC25. Therefore, we also investigated the virulence of the deletion mutant carrying a transgene encoding a version of ArPEC25 with a nuclear export signal (NES). The resulting strain, *Δarpec*25/*ArPEC25*-*NES*, was compromised in its virulence against chickpea to the same extent as the deletion mutant (Figure 1D and Supplemental Figure S1C). Statistical analyses of bioassay data from three independent experiments indicated significant differences in pathogenicity symptoms (in terms of lesion number and diameter) between the WT and *Δarpec25* (Figure 1, E and F). Moreover, *ArPEC25* transcript levels increased 500-fold and 250-fold at 12 and 24 h post inoculation (hpi), respectively, in chickpea plants infected with WT *A. rabiei* compared to mock-treated plants, as determined by real-time PCR (Figure 1C). *ArPEC25* transcript levels also rose 10-fold in *A. rabiei* mycelia 3 h after menadione treatment, which is a chemical agent use to mimic host-induced oxidative stress (Figure 1B). We also confirmed that the reduced virulence of the *ArΔpec25* strain on chickpea is not a defect in initial germination or growth of fungal mycelia, as shown by microscopy (Supplemental Figure S1E). These results suggest that ArPEC25 is important for the full virulence of *A. rabiei*, possibly by functioning as an effector whose nuclear localization is a prerequisite for pathogenicity.

### The effector ArPEC25 is secreted

If ArPEC25 is an effector protein, it should be readily secreted at the site of infection, either by the conventional or nonconventional pathways. The conventional pathway involves a cleavable N-terminal signal peptide (SP) that is processed in the endoplasmic reticulum (ER) before secretion (Liu et al., 2014). To explore ArPEC25 localization in *A. rabiei* invasive hyphae, we tagged the effector with yellow fluorescent protein (YFP) at its C terminus (ArPEC25-YFP) and introduced the corresponding construct into the fungus. Visualization by confocal microscopy detected the effector throughout the hyphae in the form of punctate structures that readily redistributed around the hyphal periphery upon H_2_O_2_ treatment, suggesting the probable site of ArPEC25 secretion (Supplemental Figure S6C).

To validate the secretory nature of ArPEC25, we employed a secretion trap assay in yeast (*Saccharomyces cerevisiae*) (Gao et al., 2019; Jiang et al., 2020). Accordingly, we generated the constructs *ArPEC25-Suc2* and *ArPEC*25*ΔSP-Suc2* by cloning the coding sequence for full-length and SP-truncated *ArPEC25* into the pYST1 vector, followed by transformation into the yeast *suc*2 mutant strain, which lacks the secreted invertase Suc2p (Supplemental Figure S8A). The resulting *ArPEC25-Suc*2 transformants grew on sucrose agar plates, which can only support the growth of yeast cells that secrete invertase, while transformants harboring the empty vector control (EV) or the *ArPEC*25*ΔSP* construct did not (Figure 3A). We also confirmed the secretion of ArPEC25 by testing for invertase-mediated reduction of 2,3,5-triphenyltetrazolium chloride (TTC) into the insoluble red product triphenylformazan, which was only observed in yeast transformants harboring *ArPEC*25 (Figure 3B).

Next, we performed an immunoblot experiment to detect the secreted ArPEC25 in culture filtrates (CF) from *A. rabiei* overexpression (OE) transformants expressing a C-terminally Flag-tagged *ArPEC25* under the control of the constitutive *Glyceraldehyde-3-phosphate dehydrogenase* (*GPDA*) promoter. Indeed, we obtained a strong immunoblot signal from the CF and fungal hyphae of the OE transformants using an anti-Flag antibody (Figure 3C and Supplemental Figure S5A). We immunopurified the secreted effector in the CF using an anti-Flag antibody and subjected the protein fraction to liquid chromatography-tandem mass spectrometry (LC-MS/MS) analysis to confirm the identity of the protein as ArPEC25 (Supplemental Figure S5, B and C). Several peptides matched the ArPEC25 protein sequence; the MS data also revealed that the N-terminal 28-aa SP is cleaved before secretion (Figure 3, D and E).

The translocation of several *Plasmodium* proteins, such as histidine-rich protein II (HRPII), knob associated histidine-rich protein (KAHRP), and glycophorin binding protein 130 (GBP130), from the parasite to host erythrocytes is tightly regulated by the conserved PEXEL motif RxLxE/D/Q (Boddey et al., 2009). To determine if the *A. rabiei* PEXEL sequence RTLND performs a similar function, we generated various mutant constructs where the first, third, and/or fifth conserved residues were mutated, resulting in the constructs *ArPEC25-Flag-YFP*, *ArPEC25^ATAND^-Flag-YFP*, and *ArPEC25^RTLNA^-Flag-YFP* (Supplemental Figure S4A). We confirmed the accumulation of the protein encoded by the chimeric constructs by immunoblot with an anti-Flag antibody and through confocal microscopy for the YFP signal (Supplemental Figure S4, B and C). Surprisingly, we detected bands of the expected size for both the wild-type and mutant proteins from the CF of axenically grown transformants (Figure 2B). We used the non-secretory fungal protein Old Yellow Enzyme 6 (ArOye6) from *A. rabiei* as a control for potential contamination with cytosolic proteins in the CF. In contrast to *Plasmodium* effector proteins whose secretion requires cleavage of the conserved PEXEL motif at the leucine (L) residue, ArPEC25 LC-MS/MS spectra indicated that the secreted effector purified from the CF retains an intact PEXEL motif, suggesting that cleavage of the PEXEL motif is not a requirement for effector secretion from the fungus (Supplemental Figure S5D). Collectively, these results confirm that ArPEC25 is secreted through the conventional pathway and show that unlike *Plasmodium* effectors, the PEXEL sequence of ArPEC25 is not cleaved during effector secretion from the fungus.

**Figure 2.**
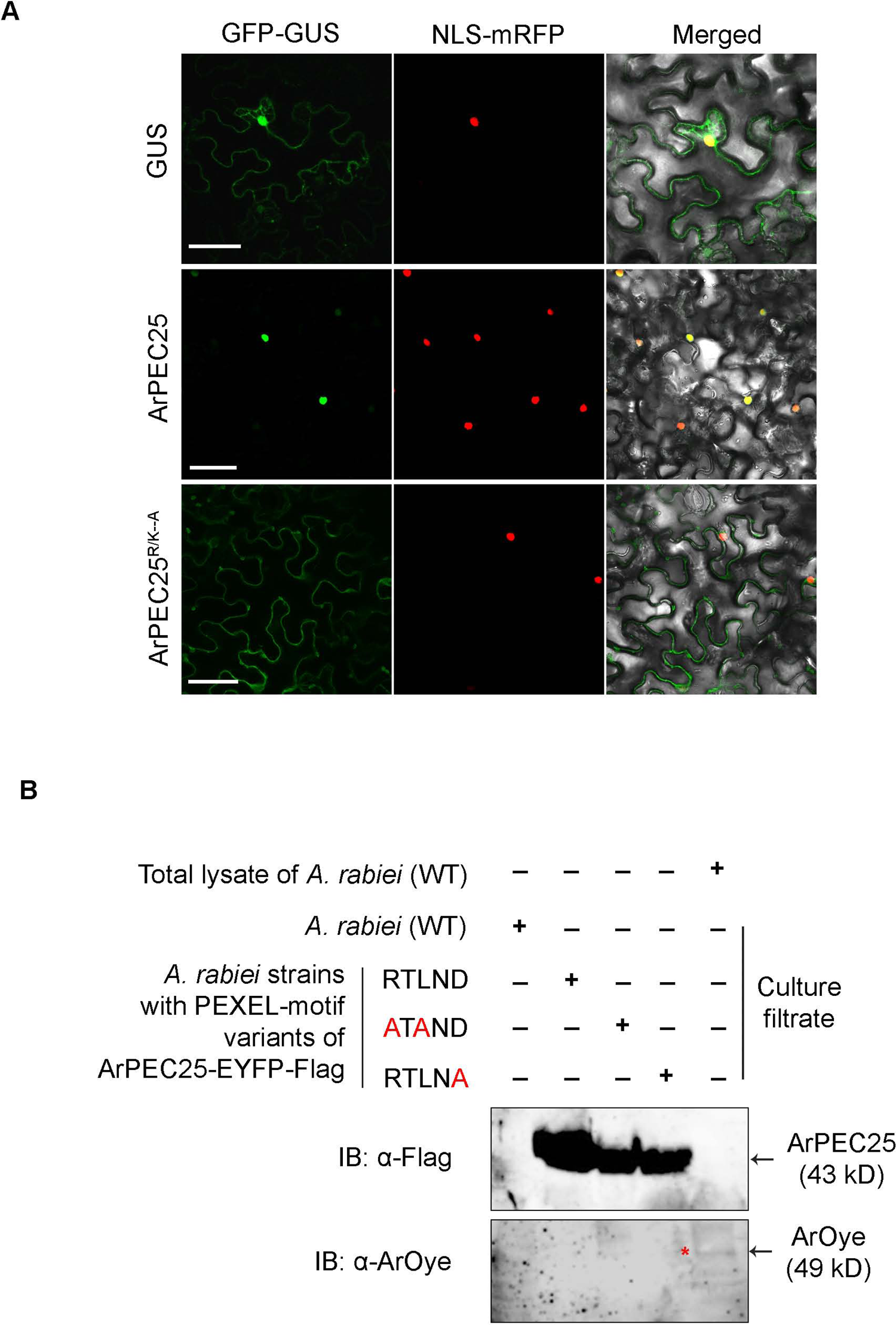
ArPEC25 localizes to plant cell nucleus and PEXEL sequence is unprocessed. **(A)** Localization study of ArPEC25 in *Nicotiana benthamiana* leaves. Agrobacterium strain containing indicated plasmids were co-infiltrated in leaf cells and fluorescence signal was detected 48 hpi. ArPEC25ΔSP-GFP-GUS localizes to nucleus (middle panel) whereas, ArPEC25 with mutated NLS, ArPEC25ΔSPAAAAAAA-GFP-GUS missed the nuclear localization (lower panel). Upper panel shows the localization of empty vector control, GFP-GUS in the cell. Bar, 50 µm. **(B)** PEXEL impart no role in effector secretion from *A. rabiei*. Various mutant translationally fused to Flag-YFP at C-terminal; ArPEC25 ^RTLND^-Flag-YFP, ArPEC25^ATAND^-Flag-YFP and ArPEC25^RTLNA^-Flag-YFP were expressed in wild type *A. rabiei*. Processed culture filtrate (CF) was analyzed for ArPEC25 secretion through immunoblots using anti-Flag Ab (upper panel) and ArOYE6 antibody (lower panel). ArPEC25 bands were detected in all transformants. CF and total lysate of wild type *A. rabiei* was used as control. Anti-ArOYE6 antibody was used for cell lysate contamination. Red color asterisk represents ArOYE6 band in total fungal lysate. ‘+’ and ‘–’ shows presence and absence, respectively.

**Figure 3.**
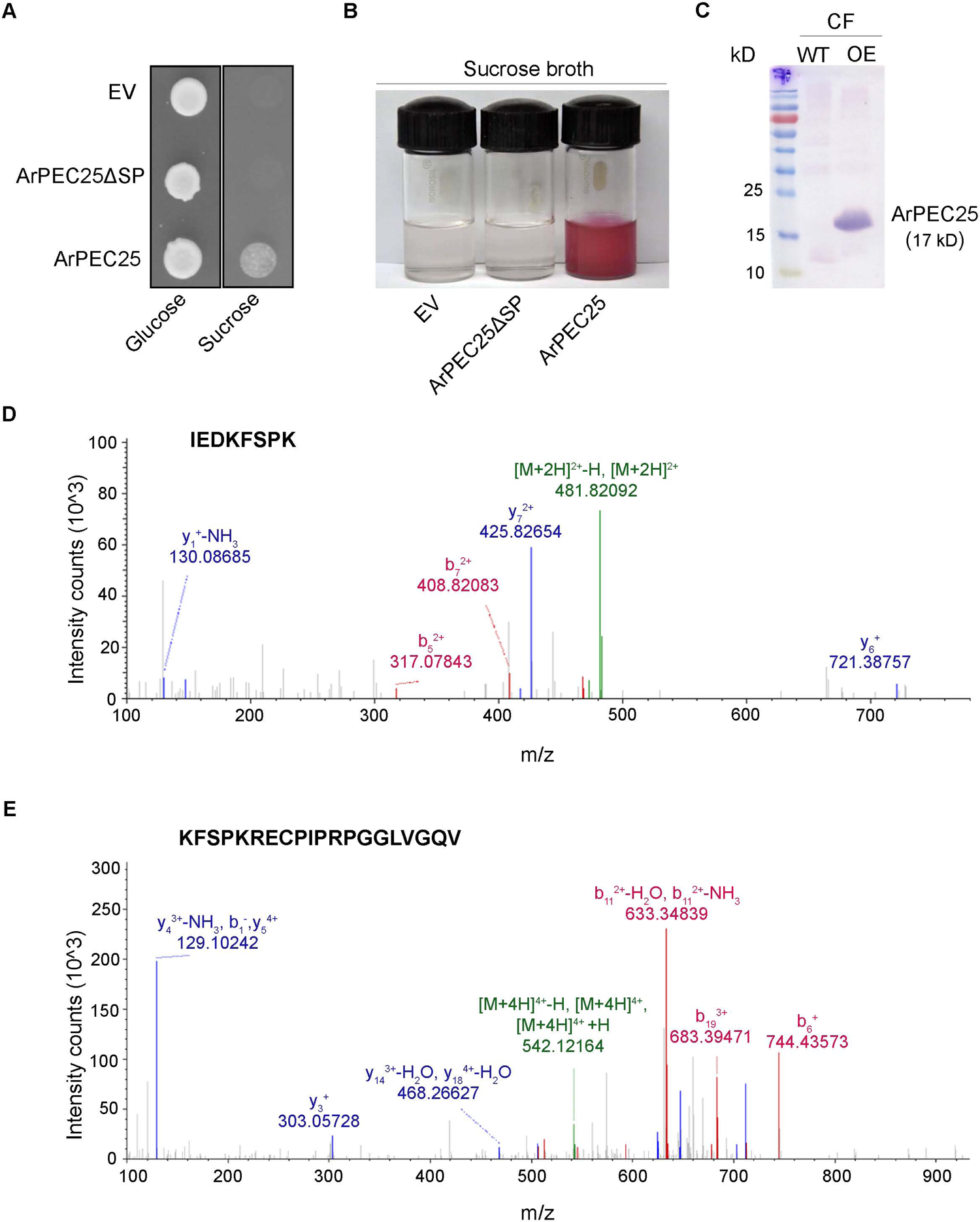
ArPEC25 is secretory in nature and its signal peptide (SP) is cleaved before secretion. **(A)** and **(B)** ArPEC25 contains a functional signal peptide as revealed in yeast secretion trap (YST) system. **(A)** The *suc2* mutant containing the indicated plasmids were grown on indicated selection medium. Growth of transformed yeast cells on sucrose medium indicates secretion of ArPEC25. **(B)** TTC (triphenyltetrazolium chloride) assay validate ArPEC25 secretion. The red color CF of *suc2* yeast cells expressing full length ArPEC25 suggests its secretion in the CF. The *suc2* strain carrying ArPEC25ΔSP and empty vector (EV) served as a control. **(C)** Effector ArPEC25 is detected in culture filtrate. The PDB culture of *A. rabiei* (WT) and over expression (OE) transformant expressing ArPEC25-Flag was processed and secretion of the protein in CF was detected with anti-Flag antibody. **(D) and (E)** Mass spectrometry of secreted ArPEC25. Processed CF of OE used in **(C)** was used for immuno purification of secreted ArPEC25. Purified protein was analyzed for the cleavage of N-terminal SP using LC-MS/MS. The peptide fragments in **(D)** and **(E)** correspond to ArPEC25. Absence of peptide fragment with signal sequence indicates its cleavage.

### ArPEC25 localizes to the nucleus in planta

ArPEC25 was predicted to localize to the host nucleus (Figure 1A), prompting us to functionally test this prediction with a construct expressing a translational fusion between GFP, β-GLUCURONIDASE (GUS), and ArPEC25 lacking the SP, yielding the construct *ArPEC25ΔSP-GFP-GUS*. The construct was transiently co-infiltrated into *Nicotiana benthamiana* leaves with the nuclear marker control construct *NLS-RFP* (red fluorescent protein) via Agrobacterium (*Agrobacterium tumefaciens*)-mediated infiltration. We observed GFP fluorescence exclusively in the nucleus (middle and upper panel of Figure 2A and Supplemental Figure S6A, respectively), suggesting that ArPEC25 is a nuclear effector.

Next, we mutated all residues in the predicted ArPEC25 NLS to alanine (A) in the *ArPEC25ΔSP-GFP-*GUS construct, resulting in *ArPEC25ΔSP^AAAAAAA^-GFP-GUS* for transient co-infiltration. The mutations to alanine abolished the accumulation of the fusion protein in the nucleus, showing a cytosolic pattern instead (lower panel of Figure 2A and Supplemental Figure S6A). These results indicate that the sequence RKRRRRR in ArPEC25 is a functional NLS that is essential for the nuclear localization of the effector during pathogenesis.

### ArPEC25 interacts with the chickpea transcription factor CaβLIM1a

Pathogen-delivered effector molecules physically interact with an array of host proteins and interfere with their normal functions of signaling, transcription, or physiological processes to render the host plant susceptible to infection. Identification of these targets is a promising approach to elucidate effector function during plant infection (Pogorelko et al., 2016). Accordingly, we used yeast two-hybrid (Y2H) screening based on the split-ubiquitin system and a bait construct harboring the *ArPEC25* (without SP) coding sequence and screened a cDNA library prepared from RNA extracted from chickpea tissues infected with *A. rabiei*. We obtained eight putative ArPEC25-interacting clones belonging to the LIM (Lin 11, Isl-1, and Mec-3) and TCP (TEOSINTE BRANCHED 1, CYCLOIDEA, and PROLIFERATING CELL NUCLEAR ANTIGEN FACTOR1/2) families of transcription factors (Supplemental Table S2). The chickpea genome encodes nine proteins with similarity to the eukaryotic lineage-specific subfamily of two LIM domain (2LIM) proteins (Srivastava and Verma, 2015). Of those, *CaβLIM1a* was mostly expressed in stems and exhibited a steep rise in its transcript levels immediately upon *A. rabiei* infection that remained high for 12 hpi and declined thereafter (Srivastava and Verma, 2015). CaβLIM1a was also the only member of the chickpea 2LIM family that is predicted to have transactivator activity. We therefore selected CaβLIM1a for further characterization.

We generated a prey construct consisting of the full-length *CaβLIM1a* coding sequence to confirm its interaction with ArPEC25 in the yeast split-ubiquitin assay (Figure 4A and Supplemental Figure S7A). We independently validated their interaction in plant cells by bimolecular fluorescence complementation (BiFC) assay by tagging CaβLIM1a with the N-terminal half of the yellow fluorescent protein Venus and ArPEC25 with the C-terminal half of Venus, resulting in the constructs *CaβLIM1a-NVenus* and *ArPEC25-CVenus* (Figure 4C). We transiently co-infiltrated both the constructs with the *NLS-RFP* marker into *N. benthamiana* leaves by Agrobacterium-mediated infiltration. We observed YFP signals in the nucleus of leaf epidermal cells, but not in the cytoplasm (lower panel, Figure 4D), indicating that ArPEC25 and CaβLIM1a interact in the plant cell nucleus.

**Figure 4.**
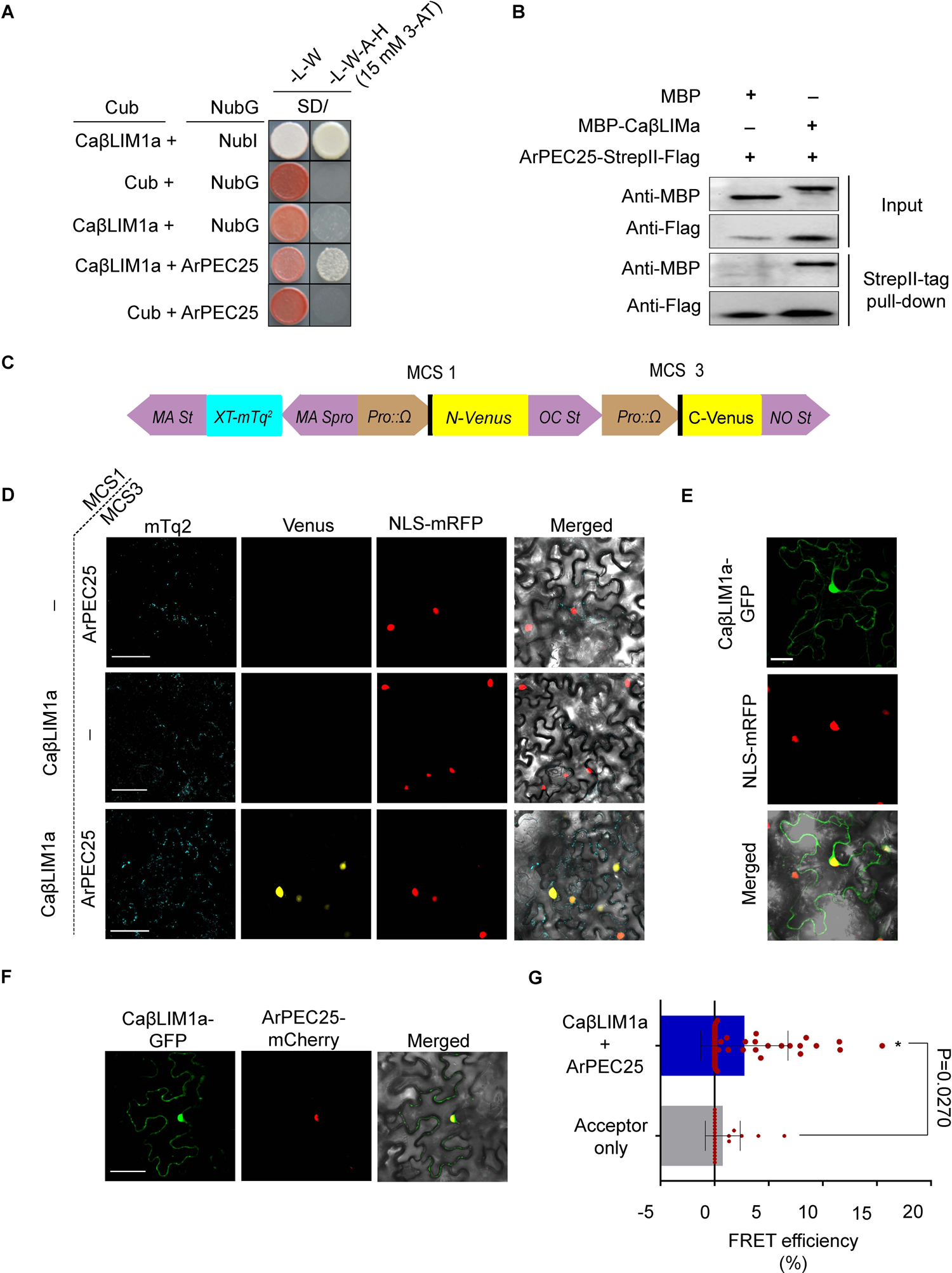
ArPEC25 interacts and co-localizes with chickpea transcription factor CaβLIM1a. **(A)** Split ubiquitin-based Yeast Two-hybrid assay indicate interaction of ArPEC25 and CaβLIM1a. Yeast cells containing the indicated plasmids were grown on indicted selective medium. The Large T and Δp53 are positive controls while pAI-Alg5 and pDL2-Alg5 are bait expression and autoactivation control, respectively. 3-AT, 3-Amino-1,2,4-triazole; SD(-LW), synthetic dextrose (-Leu, -Trp) medium; SD(-LWAH), synthetic dextrose (-Trp,-Leu,-His,-Ade) medium. **(B)** *In vitro* pull-down assay. Purified MBP, MBP-CaβLIM1a and ArPEC25-Strep II-Flag were used for the pull-down assay. Input blots show presence of purified proteins with indicated anti body. Pull-down blot shows the interaction between the proteins with anti-MBP antibody. MBP alone was taken as control. ‘+’ and ‘–’ shows presence and absence, respectively. **(C)** Schematic diagram of vector used in BiFC assay. The map shows the site of cloning of CaβLIM1a and ArPEC25 used for the BiFC assay in **(D)**. **(D)** ArPEC25 interacts with CaβLIM1a as revealed by BiFC assay in *N. benthamiana*. Translational fusion of CaβLIM1a-NVenus and ArPEC25-CVenus was transiently expressed in leaf cells. Detection of YFP fluorescence indicates an interaction between the two proteins. Detection of mTurquoise2 (mTq2) signal indicates successful transient expression in the leaf cells. Bar, 5 µm. **(E)** Sub-cellular localization of CaβLIM1a in *N. benthamiana*. CaβLIM1a-GFP was transiently co-expressed with nuclear marker (NLS-RFP) in leaf cells. The protein localizes to nucleus as revealed by merged image. Bar, 5 µm. **(F)** Co-localization of ArPEC25 and CaβLIM1a in *N. benthamiana* Indicated CaβLIM1a-GFP and ArPEC25-mCherry were co-expressed in leaf cells. Merged image indicates co-localization of the two protein in nucleus. Bar, 5 µm. **(G)** ArPEC25 interacts with CaβLIM1a as revealed in FRET. The donor CaβLIM1a-GFP and the acceptor ArPEC25-mCherry were transiently co-expressed in *N. benthamiana* leaves. The FRET was analyzed 48 hpi using confocal microscopy. Error bar represents the ± SD from three independent biologicals replicates (n=3). The acceptor only represents ArPEC25-mCherry and GFP. Statistically significant differences were calculated by unpaired one tailed t-test.

We further explored their physical interaction in vivo using Förster resonance energy transfer-acceptor photobleaching (FRET-APB) by constructing translational fusion constructs between CaβLIM1a and the FRET donor GFP at its C terminus, while ArPEC25 was fused to the FRET acceptor mCherry at its C terminus. The resulting constructs were then transiently co-infiltrated in *N. benthamiana* leaves (Supplemental Figure S7B). We measured strong FRET efficiency (with a mean value of 2.745 ± 0.62) compared to controls (mean value of 0.76 ± 0.33), in agreement with an interaction between CaβLIM1a and ArPEC25 (Figure 4G).

We also tested the interaction between ArPEC25 and CaβLIM1a by in vitro pull-down assays using recombinant maltose-binding protein (MBP)-CaβLIM1a and streptavidin II (Step II)-Flag-ArPEC25. Recombinant MBP-CaβLIM1a was pulled down with ArPEC25-Strep II-Flag in vitro, but not MBP alone, as determined by immunoblots (Figure 4B), supporting the interaction of the two proteins. The physical association between the two proteins was also validated by the detection of recombinant protein MBP-CaβLIM1a-6xHis and ArPEC25-6xHis with anti-His antibody (Figure 7B) in the same elution fraction derived from gel filtration chromatography assay.

### CaβLIM1a colocalizes with ArPEC25 in the nucleus

Plant LIM transcription factors localize to the cytoplasm, the nucleus, or both compartments to regulate transcription (Hoffmann et al., 2017; Yang et al., 2019). To investigate the subcellular location of CaβLIM1a, we generated a construct placing the full-length coding sequence of *CaβLIM1a* upstream and in-frame with that of *GFP*. We then transiently co-infiltrated the resulting construct *CaβLIM1a-GFP* and the nuclear control *NLS-RFP* into *N. benthamiana* leaves. The CaβLIM1a-GFP fusion protein was localized to both the cytosol and the nucleus, as the GFP signal overlapped with that of RFP (Figure 4E). We also transiently co-infiltrated *N. benthamiana* leaves with *CaβLIM1a-GFP* and *ArPEC25-mCherry* constructs to investigate the co-localization of the two proteins in planta. Indeed, CaβLIM1a colocalized with the effector exclusively in the plant nucleus (Figure 4F).

### CaβLIM1a is an activator that may modulate the expression of *CaPAL1*

Plant LIM transcription factors have been described as having transcriptional activator or repressor properties in many plant species (Kim and Hwang, 2014; Li et al., 2018). Therefore, we attempted to decipher the transcriptional function of CaβLIM1a and other 2LIM proteins from chickpea (Ca2LIMs) isolated using the split-ubiquitin assay. We conducted a transcriptional activator assay, whereby the full-length coding sequences of *Ca2LIM*s fused to the DNA-binding domain (DB) or the yeast GAL4 protein were expressed in yeast strain Y2H Gold under the control of the *ALCOHOL DEHYDROGENASE 1* (*ADH1*) promoter. Of the six LIM proteins tested in this assay (CaWLIM1a, CaWLIM1b, CaβLIM1a, CaβLIM1b, CaWLIM2, and CaδLIM2), only CaβLIM1a activated the transcription of reporter genes driven by GAL4 regulatory sequences, as evidenced by the growth of yeast colonies on selective growth medium (Figure 5A and Supplemental Figure S9A), suggesting that CaβLIM1a is a transcriptional activator. None of the reporter genes showed any expression in yeast colonies expressing *GAL4 DB* alone. A repressor assay in yeast suggested that none of the Ca2LIMs are transcriptional repressors, as they failed to block the activation of reporter genes driven by the herpes simplex virus virion protein (VP16) regulatory sequences when fused to the activator domain of the strong transactivator VP16 (Supplemental Figure S9C).

**Figure 5.**
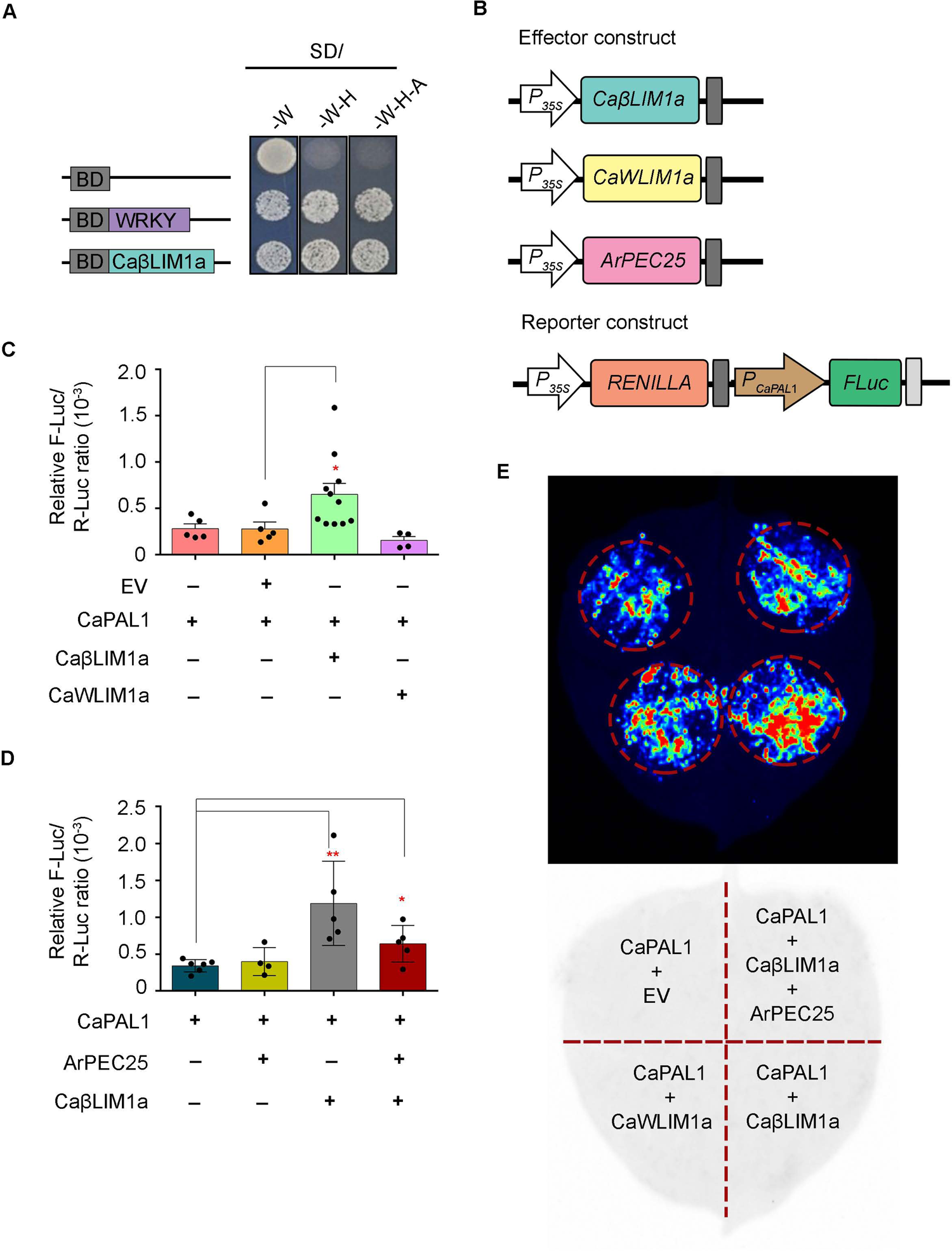
Transcription factor CaβLIM1a is activator. **(A)** Transactivation activity assay of CaβLIM1a in yeast system. Yeast strain containing indicated plasmids were grown on selective medium. CaβLIM1 activated the expression of reporter gene as revealed by the growth of cells on selective medium. WRKY50 and empty vector (BD) served as positive and negative control, respectively. SD/(-W); synthetic dextrose (-Trp) medium, SD/(-W-H); synthetic dextrose (-Trp, -His) and SD/(-W-H-A), synthetic dextrose (-Trp, -His, -Ade). **(B), (C), (D) and (E)** Transient luciferase (LUC) reporter assay in *N. benthamiana* leaves. **(B)** Schematics of different reporter and effector constructs used in **(C)**, **(D)** and **(E)**. Reporter construct contain *CaPAL*1 promoter upstream to firefly luciferase (*FLuc*) gene. **(C)** Quantitative analysis of reporter assay in *N. benthamiana* leaves. Each construct shown in **(B)** were infiltrated in combination and relative LUC activity was measured. Expression data represent mean ± SD of 5 to 11 biological replicates (n=5 and n=11). Statistically significant differences were calculated by unpaired one tailed t-test; * P ≤ 0.0324. “+” and “-” indicate presence and absence, respectively. **(D)** ArPEC25 inhibits CaβLIM1a transactivation function as revealed by dual-LUC assay in *N. benthamiana*. Constructs in **(B)** were transiently expressed in combination in leaves for relative LUC activity. Data represent mean ± SD of 5 to 6 independent experiment (n=5 and n=6). Statistically significant differences were calculated by unpaired one tailed t-test; * P ≤ 0.0106, ** P ≤ 0.0055. “+” and “-” indicate presence and absence, respectively. **(E)** Qualitative analysis of reporter assay in *N. benthamiana* leaves. The reporter and effector constructs shown in **(B)** were infiltrated in combination and image taken 48 hpi. Upper panel shows the intensity of the expression of luciferase and lower panel shows the scheme of infiltration.

The gene encoding the enzyme phenylalanine ammonia lyase (PAL) shows differential regulation during biotic stress such as fungal attacks (Zhang et al., 2017). The chickpea genome contains four *PAL* genes: *CaPAL1* (LOC101507594), *CaPAL2* (LOC101509831), *CaPAL3* (LOC101496077), and *CaPAL4* (LOC101493062). We investigated the differential expression of *PAL* genes in chickpea plants infected with *A. rabiei* and mock-infected plants. Only *CaPAL1* displayed a strong biphasic induction of its transcript levels, as seen by real-time PCR (Supplemental Figure S9D). These results suggest that *PAL1* transcription is activated upon *A. rabiei* infection, possibly via CaβLIM1a transactivator function.

### CaβLIM1a binds to the *CaPAL1* promoter sequence and positively modulates *CaPAL1* **expression**

Plant LIM transcription factors have been reported to regulate the phenylpropanoid biosynthetic pathway by binding to the *PAL-box* element, with the consensus sequence CCA(C/A)C(A/T)A(C/A)C(C/T)CC (Kawaoka et al., 2000). As *CaPAL1* transcript levels were strongly upregulated in response to *A. rabiei* infection, we hypothesized that CaβLIM1a may be involved in this transcriptional response. To test this hypothesis, we searched the promoter sequences of all four *CaPAL* genes for a *PAL-box*. Notably, we identified two *PAL-box* elements (*PAL-box*1 and *PAL-box*2) near one another in the *CaPAL1* promoter, but not in the other *CaPAL* promoters (Figure 6A). We then performed a yeast one-hybrid assay (Y1H) to assess the binding of CaβLIM1a to the *CaPAL1* promoter. We cloned a promoter fragment extending 737 bp upstream of the transcription start site (TSS) of *CaPAL1*, which included *PAL-box1* and *PAL-box2*, and placed it upstream of the *AUR1-C* reporter gene in the pABAi vector, which confers resistance to the antibiotic Aureobasidin A (AbA) if expressed. In parallel, we cloned the full-length *CaβLIM1a* coding sequence downstream and in-frame with the GAL4 activation domain (AD) in the pGADT7 vector, driven by the yeast *ADH1* promoter. Importantly, yeast colonies harboring both the *CaPAL1pro:AUR1-C* and *AD-CaβLIM1a* constructs showed activation of the AUR1-C reporter, as demonstrated by the growth of colonies on medium containing AbA (Supplemental Figure S10A). The two other chickpea LIM proteins, CaWLIM2 and CaWLIM1a, also bound to the same promoter fragment (Supplemental Figure S10A). We selected only CaβLIM1a for further characterization because CaWLIM2 and CaWLIM1a lacked transactivator or transrepressor activity (Supplemental Figure S9, A and C). Next, we investigated the functional relevance of each *PAL-box* in the *CaPAL1* promoter by generating promoter constructs lacking one or both of the *PAL-box* elements: *CaPAL1^ΔBox1^*, *CaPAL1^ΔBox2^*, and *CaPAL1^ΔBox1 ΔBox2^* (Supplemental Figure S8D). Both *PAL-boxes* were crucial for binding by CaβLIM1a in Y1H assays, as none of the *PAL-box* deletion constructs sustained yeast growth on medium containing AbA (Figure 6C).

**Figure 6.**
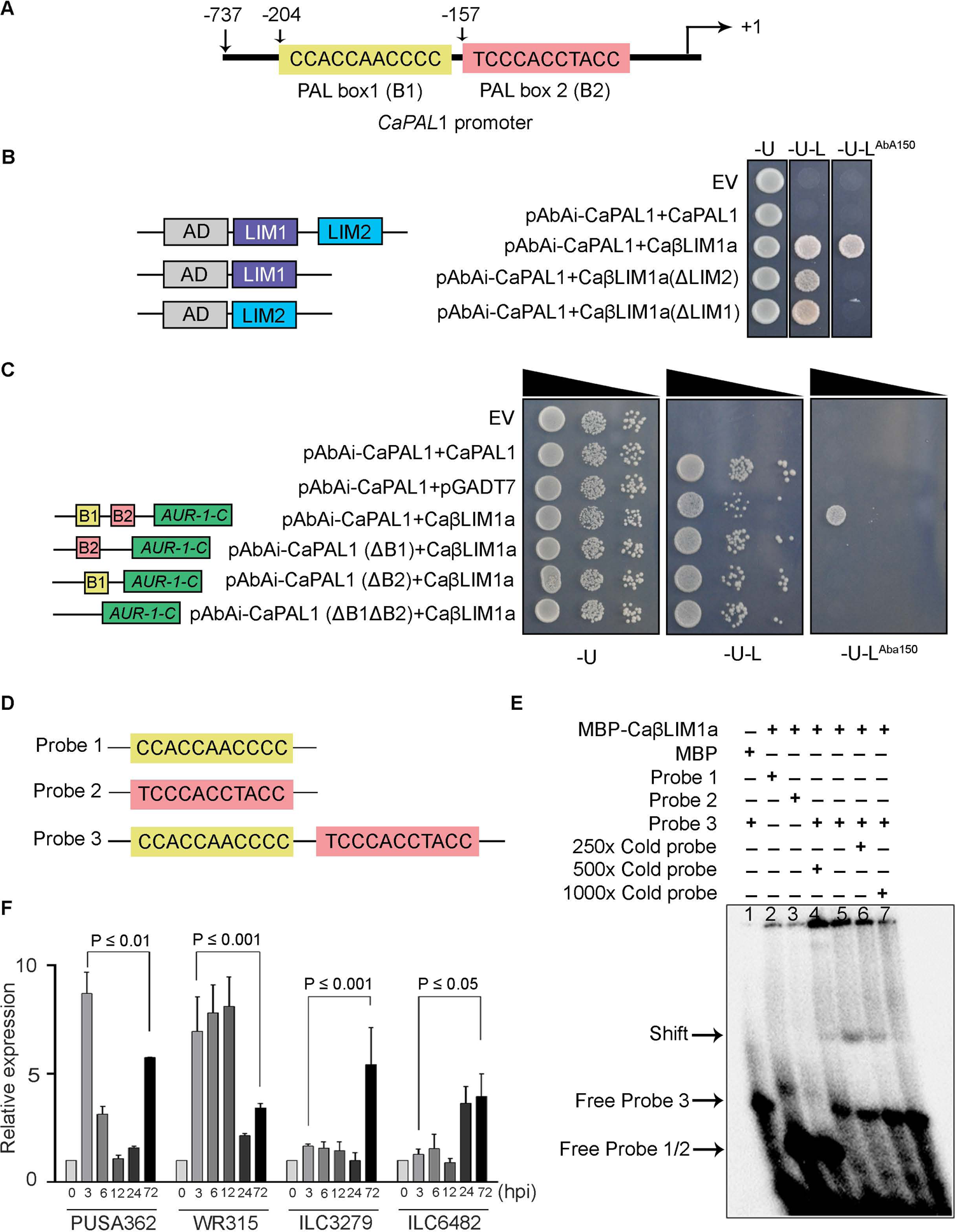
CaβLIM1a bind on the promoter of *CaPAL*1 gene. **(A)** Schematic diagram of *CaPAL*1 promoter. The number on *PAL-box*1 (B1) and *PAL-box*2 (B2) of *CaPAL*1 promoter in the schematics represents box position. **(B)** CaβLIM1a bind to the promoter in Y1H assay. Yeast strain containing indicated plasmids were grown on selective medium. Growth of cells on medium supplemented with 150 ng/ml of AbA indicates DNA protein interaction. LIM domain deletion constructs failed to interact with DNA. SD/(-U) synthetic dextrose (-Ura), SD/(-U-L), synthetic dextrose (-Ura-Leu), SD/(-U-L^AbA150^), synthetic dextrose (-Ura-Leu supplemented with 150 ng/ml aureobasidin A). **(C)** Importance of PAL boxes in DNA protein interaction. Yeast cells containing indicated plasmids were grown on selective medium. Growth of cells in AbA supplemented medium suggests interaction. PAL-box deletion constructs (*CaPAL*1*^ΔB^*^1^, *CaPAL*1*^ΔB^*^2^ and *CaPAL1^ΔBox1 ΔBox2^*) failed to interact with CaβLIM1a. SD/(-U), synthetic dextrose (-Ura), SD/(-U-L), synthetic dextrose (-Ura-Leu), SD/(-U-L^AbA150^), synthetic dextrose (-Ura-Leu supplemented with 150 ng/ml aureobasidin A). **(D)** and **(E)** EMSA using MBP-CaβLIM1a and *PAL-box*1 and *PAL-box*2 sequence as probes. **(D)** Synthesized probe 1, probe 2, and probe 3 used in the assay. **(E)** EMSA experiment using probes 3 in **(D)** and purified MBP-CaβLIM1a. Binding between CaβLIM1a and probe 3 is detected on the membrane. The cold probe in the indicated concentration range were used as competitor. “+” and “-” indicate presence and absence, respectively. Purified MBP served as a control. **(F)** Expression analysis of *CaPAL*1 in chickpea susceptible and resistant variety infected with A. *rabiei*. mRNA was extracted from susceptible variety, Pusa362 and WR315 and resistant variety, ILC 3279 and ILC 6482 at indicated time points and relative expression of *CaPAL1* was analyzed. Error bar represents ± SD from two independent biologicals replicates each with three technical replicates. Transcript abundance was normalized to that of the expression of the control chickpea *CaβTub*.

To further evaluate the importance of *PAL-box1* and *PAL-box2* in the binding of CaβLIM1a to DNA, we performed electrophoretic mobility shift assays (EMSAs) using recombinant CaβLIM1a and ArPEC25 purified from *E. coli* (Supplemental Figure S10E). We designed three radiolabeled probes: *P1* (42 bp), *P2* (41 bp), and *P3* (70 bp) containing *PAL-box1*, *PAL-box2*, and *PAL-box1*-*PAL-box2*, respectively (Figure 6D). After incubation of recombinant CaβLIM1a with the individual probes, we detected a mobility shift only for the *P3* probe following electrophoresis (Figure 6E). Binding of CaβLIM1a to the *P3* probe gradually diminished upon co-incubation with increasing excess unlabeled *P3* probe (*P3*) (Figure 6E). We failed to observe shifted radiolabeled bands when CaβLIM1a was incubated with *P1* or *P2* (Figure 6E), suggesting that both *PAL-box1* and *PAL-box2* must be present in the *CaPAL1* promoter for binding to occur.

Next, we performed a dual-luciferase assay in *N. benthamiana* leaves to confirm that CaβLIM1a can bind to the *CaPAL1* promoter in planta and to determine the type of regulation imposed by *CaPAL1* on *CaPAL1* transcription. Accordingly, we transiently co-infiltrated the *CaβLIM1a-Myc* effector construct and a reporter construct consisting of the firefly luciferase (*FLuc*) gene under the control of the *CaPAL1* promoter (*ProCaPAL*1*-FLuc*) into *N. benthamiana* leaves (Figure 5B). The luminescence signal derived from *FLuc* transcription rose above background levels only when *CaβLIM1a* was expressed in *N. benthamiana* leaves (Figure 5, C and E), confirming that the transcription factor interacts with the *CaPAL1* promoter. In addition, the observed increase in luminescence upon *CaβLIM1a* expression compared to the empty vector (EV) control suggested that CaβLIM1a activates *CaPAL1* transcription (Figure 5C). The expression of another CaLIM gene, *CaWLIM1a*, whose encoded protein lacks intrinsic activator or repressor activity, similarly did not induce *CaPAL1* transcription (Figure 5C). Importantly, all effector constructs resulted in the accumulation of their respective proteins in the infiltrated leaves, as determined by immunoblotting with an anti-Myc antibody (Supplemental Figure S9B).

The LIM1 and LIM2 domains of the tobacco (*Nicotiana tabacum*) transcription factor NtLIM1 have been shown to bind to promoter sequences independently (Kawaoka et al., 2000). We thus next investigated whether CaβLIM1a behaved similarly by generating truncated constructs expressing versions of CaβLIM1a lacking either the LIM1 domain (CaβLIM1a^ΔLIM1^) or the LIM2 domain (CaβLIM1a^ΔLIM2^) (Supplemental Figure S8C). We then tested these mutants by Y1H assay for binding to the *CaPAL1* promoter. Unlike NtLIM1, CaβLIM1a appeared to require both the LIM1 and LIM2 domains to bind to the *CaPAL1* promoter, as neither CaβLIM1a^ΔLIM1^ nor CaβLIM1a^ΔLIM2^ sustained yeast growth on medium containing AbA (Figure 6B). This inability to activate the AUR1-C reporter gene was not due to protein instability, as an immunoblot assay on yeast cell extracts with an anti-HA antibody detected proteins of the expected molecular weights (Supplemental Figure S10D).

### ArPEC25 facilitates virulence in chickpea by inhibiting CaβLIM1a function

Secreted effectors modulate host immunity by binding to or enzymatically altering the function of host molecules such as NAC-type transcription factors (Yuan et al., 2019b). We speculated that ArPEC25 had a similar mode of action in the establishment of AB disease in chickpea. To test this hypothesis, we first investigated the effect of recombinant ArPEC25 on the promoter binding ability of CaβLIM1a in EMSAs using the probe *P3*. Indeed, the band corresponding to the *P*3-CaβLIM1a complex gradually disappeared with increasing concentrations of ArPEC25 (1x, 4x, 8x, and 10x) (Figure 7A), suggesting that the effector interferes with the binding of CaβLIM1a to DNA. We also repeated the dual-LUC based assay by co-infiltrating the *ProCaPAL*1*-FLuc* reporter construct with effector constructs overexpressing *ArPEC25* and *CaβLIM1a* (Figure 5B) into *N. benthamiana* leaves. The co-expression of *ArPEC25* and *CaβLIM1a* decreased FLuc activity compared to CaβLIM1a alone (Figure 5, D and E), suggesting that ArPEC25 interferes with the transactivation function of CaβLIM1a.

### The lignin biosynthetic pathway is severely compromised during infection

PAL is the regulatory enzyme that controls flux through the phenylpropanoid biosynthetic pathway and has been extensively studied in multiple plant systems in response to biotic stress such as the pathogenic fungus *S. sclerotiorum* (Ranjan et al., 2019). The expression of several genes belonging to the phenylpropanoid biosynthetic pathway have been reported to be induced in chickpea upon *A. rabiei* infection (Kavousi et al., 2009), but it remains unclear how the pathogen modulates the contents of phenolic metabolites. Our gene expression data also indicated that *CaPAL1* expression is induced at 3 hpi with wild-type *A. rabiei* in the susceptible chickpea varieties ‘Pusa362’ and ‘WR315’ but then is downregulated at 72 hpi (Figure 6F). The *CaPAL1* expression profile was markedly different in the *A. rabiei*–resistant chickpea varieties ‘ILC3279’ and ‘ILC6482’, as *CaPAL1* expression remained at basal levels during the early phase of infection (3–12 hpi), followed by a strong upregulation at 72 hpi, although not to the same extent as that seen in susceptible varieties at 3 hpi (Figure 6F). We therefore measured the levels of metabolites associated with the phenylpropanoid biosynthetic pathway of chickpea Pusa362 plants mock-infected and infected with *A. rabiei*, with a particular focus on the flavonoid and lignin biosynthetic branches. We also characterized amino acid biosynthesis of infected and mock-infected chickpea plants through ultra high-performance liquid chromatography (UH-PLC). We observed a significant reduction in the contents of the lignin biosynthetic pathway intermediates cinnamic acid, caffeic acid, syringic acid, and chlorogenic acid at 72 hpi in *A. rabiei*–infected plants relative to mock-infected plants (Figure 8, A and Supplemental Figure S12 and S13). However, most of the key intermediates in the flavonoid and amino acid biosynthetic pathways remained unaffected (Supplemental Figures S12, S13 and S14). We also looked for the specific role of ArPEC25 in suppression of lignin biosynthesis. We tested the key lignin biosynthesis intermediates that were shown to be modulated in earlier experiments using ArPEC25 RNAi transformant (#22) (Supplemental Figure S11). The metabolites related to lignin biosynthesis intermediates were found upregulated in ArPEC25 RNAi (#22) compared to wild-type infected chickpea (Figure 8, B). Thus, the lower levels of monolignol precursors in wild-type fungus-infected chickpea compared to ArPEC25 RNAi (#22) demonstrates the direct role of ArPEC25 in actively subverting host immunity by preventing new lignin biosynthesis, which would be expected to severely compromise the structural integrity of the host cell wall.

**Figure 7.**
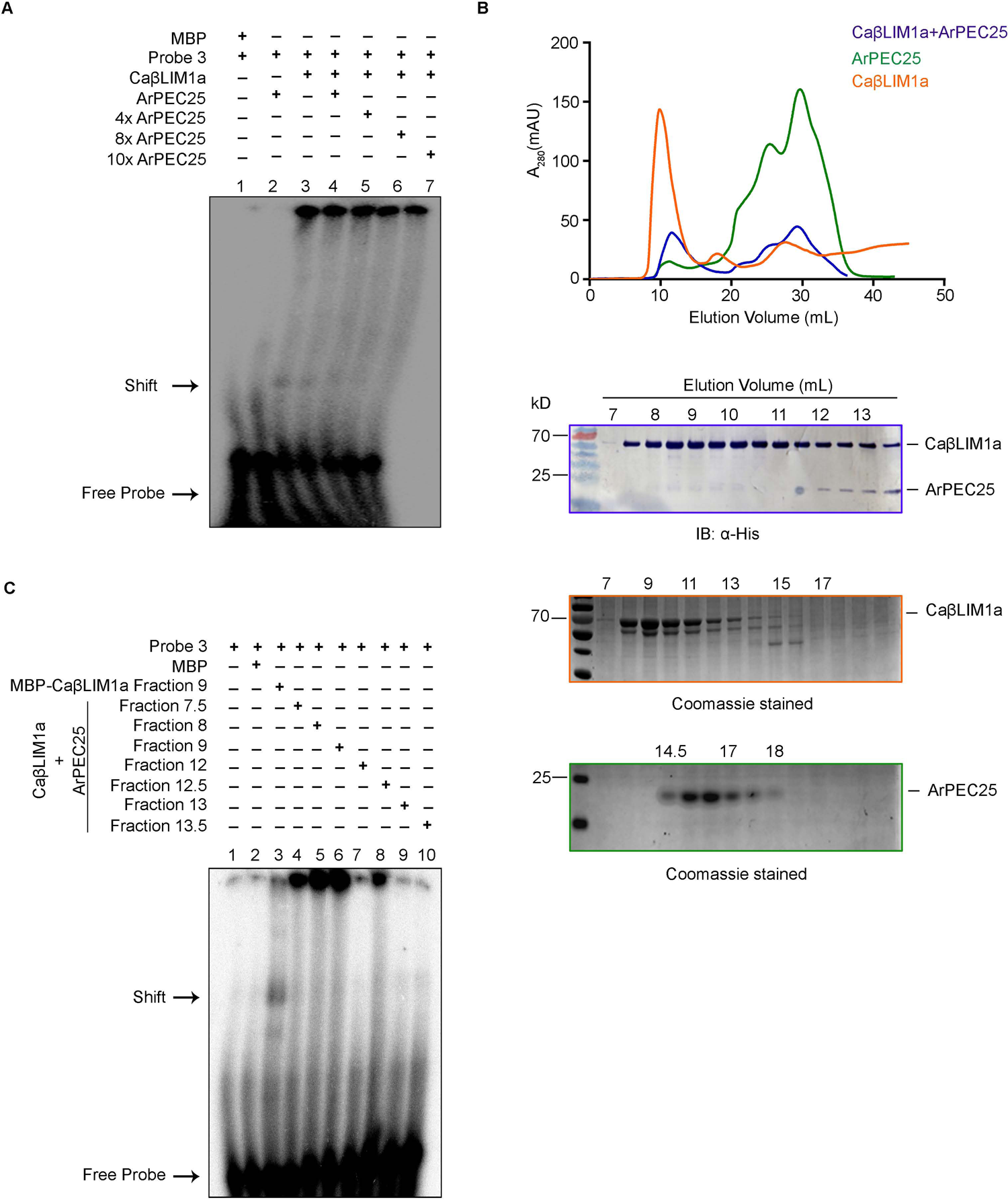
ArPEC25 inhibits CaβLIM1a binding to promoter sequence. **(A)** EMSA with purified MBP-CaβLIM1a, ArPEC25-6xHis and probe 3 in **(6D)**. Intensity of the shift shown in lane 3 gradually decreases with increasing concentration of ArPEC25 which was used as competitor. “+” and “-” indicate presence and absence, respectively. Purified MBP served as a control. **(B)** Gel filtration profile of purified CaβLIM1a and ArPEC25. Absorption peak of purified CaβLIM1a and ArPEC25 is shown with orange and green color, respectively and absorption peak with blue color represents mix of these two proteins in equal ratio. A_280_, absorbance at 280 nm and mAU refers to milli–absorbance units. Protein present in different fractions were visualized on coomassie gel and immuno blot with anti-His antibody (lower panel). **(C)** EMSA using purified MBP-CaβLIM1a, probe3 and elution fraction of mix of CaβLIM1a and ArPEC25 in **(B)**. Different protein fractions (number indicated) in **(B)** were used for EMSA. “+” and “-” indicate presence and absence, respectively. Purified MBP served as control.

**Figure 8.**
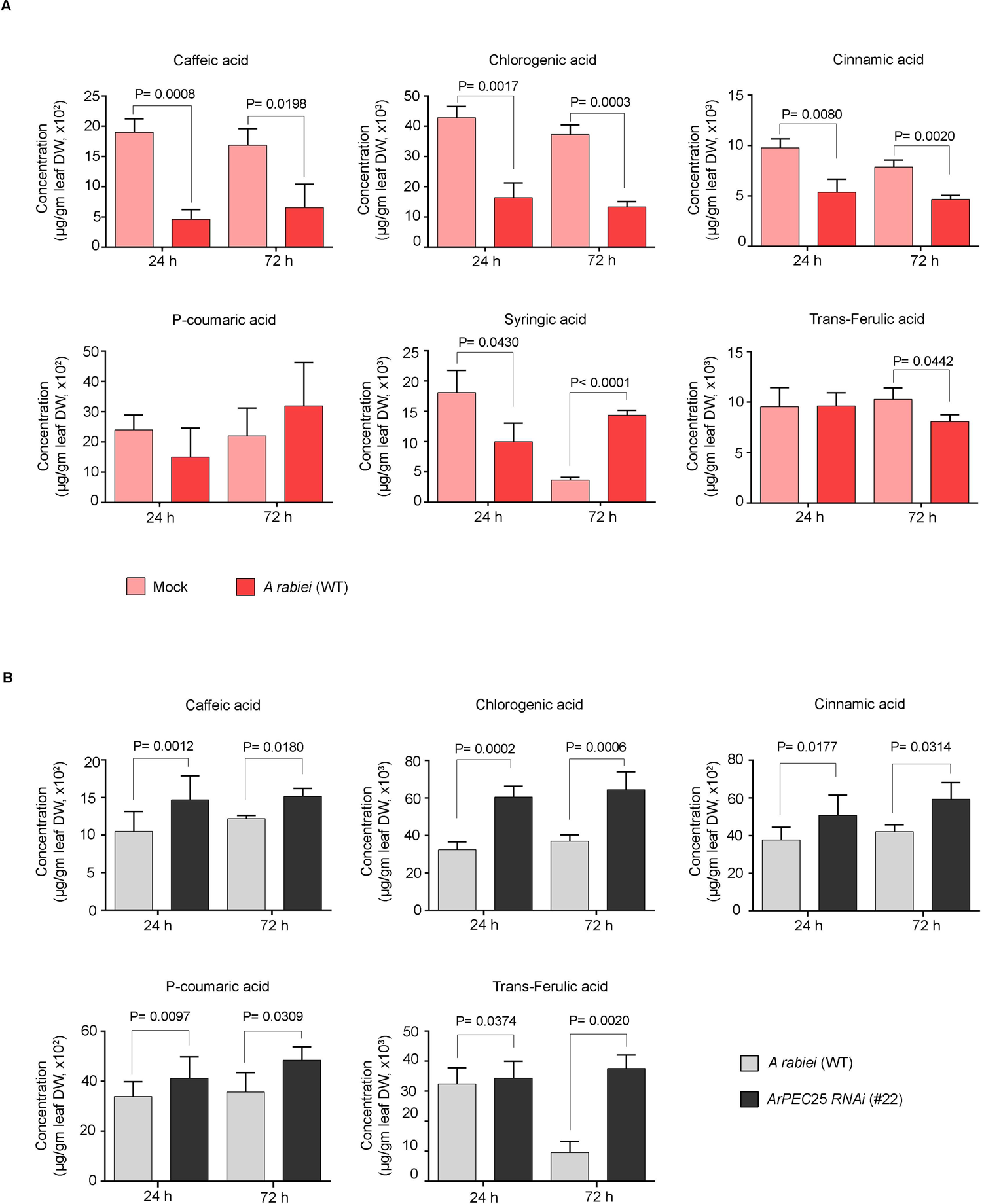
ArPEC25 negatively modulate lignin biosynthetic pathway component in infected chickpea. **(A)** and **(B)** Accumulation data are recorded for at least three independent biological replicates using UHPLC spectrum of aqueous methanolic extract of dried chickpea tissues treated with **(A)** mock and wild type *A. rabiei* and **(B)** wild type *A. rabiei* and *ArPEC25* RNAi transformants (#22). Accumulated metabolite values are plotted as fold change with respect to mock or wild type *A. rabiei*. Error bar represents ± SD value. Statistically significant differences were calculated by unpaired two-tailed in **(A)** and paired one-tailed test in **(B)**, respectively.

## Discussion

The pressures imposed by the continual evolutionary arms race between invading microbes and host plants have generated highly diversified and ever more functionally specific effectors. These effectors are generally accepted to suppress host immunity by interacting with key molecules involved in host defense to alter or disrupt their normal function. Surprisingly, for the majority of secreted effectors from fungi and oomycetes, what their host target is and how they affect its function in the context of host susceptibility remain unclear. Several studies have suggested that these targets can have diverse functions, ranging from signaling components, transcriptional regulators, metabolic enzymes, or simply products of host R-genes (Gimenez-Ibanez et al., 2014; Qin et al., 2018; Liu et al., 2014). This study on the nuclear-localized effector protein ArPEC25 revealed that its translocation to the host nucleus is crucial for pathogenesis (Figure 1D and Supplemental Figure S1C), which prompted us to hypothesize that a host nuclear protein might be its target. We identified the chickpea transcription factor CaβLIM1a as a major virulence target of ArPEC25 from a Y2H library screen. Because plant immunity relies on transcription factors (both activators and repressors) to modulate the transcription of many defense genes, it is not surprising that these transcription factors are targeted by many effectors secreted by pathogens to subvert host immunity. In fact, about 50% of all host proteins targeted by effectors participate in transcriptional regulation and in signaling (He et al., 2020). For example, the JASMONATE-ZIM-DOMAIN PROTEIN (JAZ) transcription factor is a negative modulator of JA signaling that is targeted by the MiSSP7 effector from *Laccaria bicolor* (Plett et al., 2014). Likewise, the bacterial effector XopD from *Xanthomonas campestris* subverts Arabidopsis plant immunity by repressing the function of the transcription factor MYB30, which is a positive modulator of defense genes (Canonne et al., 2011). The *P. infestans* RxLR effector Pi03192 interacts with the NAC transcription factors NAC Targeted by Phytophthora 1 (NTP1) and NTP2 of potato (*Solanum tuberosum*) and promotes virulence by preventing their accumulation in the nucleus (McLellan et al., 2013).

LIM transcription factors are characterized by two LIM domains separated by a long spacer of around 40–50 aa and have been described in various plants for their function in cytoskeleton organization and transcriptional regulation (Srivastava and Verma, 2015; Han et al., 2013; Weiskirchen and Günther, 2003; Srivastava and Verma, 2017). LIM-type transcription factors localize to the cytoplasm, the nucleus, or both (Hoffmann et al., 2017; Sala and Ampe, 2018). Cytosolic LIMs function as actin bundlers (Han et al., 2013), while nucleus-localized LIMs are generally involved in the transcriptional regulation of genes with *PAL-boxes* such as *PAL*, *CINNAMYL ALCOHOL DEHYDROGENASE* (*CAD*), and *4-COUMARATE:COA LIGASE* (*4CL*) (Kawaoka et al., 2000). We determined here that the LIM family member CaβLIM1a shows a dual localization in the cytosol and the nucleus in *N. benthamiana* leaves (Figure 4E). Similar results were reported for the cotton (*Gossypium hirsutum*) transcription factor WLIM1a, which shuttles between the nucleus and the cytosol following H_2_O_2_ treatment and also exerts a dual function in actin bundling and transcriptional activation of genes involved in the biosynthesis of lignin and lignin-like phenolic compounds, whose promoters contain *PAL-boxes* (Han et al., 2013). The secreted effector ArPEC25 is targeted exclusively to the nucleus of the host plant, where it interacts with CaβLIM1a (Figure 2A and Figure 4D). In light of its dual localization, it is possible that CaβLIM1a also functions as an actin bundler, but we have not explored this possibility here. Based on these results and our previous report on the upregulation of the expression of members of the *Ca2LIM* family of transcription factor genes upon *A. rabiei* infection (Srivastava and Verma, 2015), we propose that CaβLIM1a plays a complex role in chickpea defense through its transcriptional regulation of certain downstream genes.

To attack plant tissues, pathogens must penetrate the epidermis, either directly via stomatal openings or indirectly by employing an array of cell wall lytic enzymes. The expression of phenylpropanoid biosynthetic genes is modulated in response to biotic factors to influence the production of secondary metabolites including lignin (Zhang et al., 2017). The phenylpropanoid pathway and LIM TFs have been shown to be involved in plant-pathogen interactions in soybean, rice, cotton, and chickpea. Our data also showed an induced expression for chickpea *CaPAL1* following *A. rabiei* infection. CaβLIM1a appears to bind simultaneously to the two elements via its LIM1 and LIM2 domains in contrast to NtLIM1, for which either LIM domain is sufficient to bind to the single *PAL box* of the horseradish (*Armoracia rusticana*) *peroxidase C2* (*prxC2*) promoter (Kawaoka et al., 2000). These possible differences between chickpea and tobacco LIMs may be due to the evolutionary distance separating the two plant families.

Evidence suggests that secreted effectors target regulatory components of the phenylpropanoid pathway to subvert host responses. For example, the type-III effectors RipE1 and RipAY secreted by the necrotroph *Ralstonia solanacearum* promote infection in tobacco by enhancing the biosynthesis of SA, one of the products of the phenylpropanoid pathway (Sang et al., 2020). To support colonization and infection, effectors from necrotrophic pathogens such as ScQDO from *S. sclerotiorum* selectively hydrolyze flavonolaglycone, a product of the phenylpropanoid pathway that is normally toxic to the pathogen, to the nontoxic phloroglucinol carboxylic and phenolic acids (Chen et al., 2019). We demonstrated here that the physical interaction between the effector ArPEC25 and CaβLIM1a negatively modulates *CaPAL1* promoter activity (Figure 5, D and E). We hypothesize two possible reasons for this downregulation. First, the interaction of ArPEC25 with CaβLIM1a may disrupt oligomerization of the transcription factor, turning it into a nonfunctional protein, as transcription factors typically work as oligomers (Sayou et al., 2016). Second, ArPEC25 may interfere with the DNA binding ability of CaβLIM1a. However, gel filtration chromatography data suggested that the interaction between the two proteins has no influence on the oligomeric state of CaβLIM1a. Rather, the inability of CaβLIM1a to bind to the *PAL-box* elements of the *CaPAL1* promoter in EMSAs with increasing ArPEC25 concentrations suggests that ArPEC25 prevents the DNA binding and/or transactivation function of CaβLIM1a. Since both LIM1 and LIM2 domains are required for DNA binding by CaβLIM1a, ArPEC25 may bind to one or both domains and thus interfere with CaβLIM1a function. However, additional data are required to make a conclusive statement.

As CaβLIM1a induced *CaPAL1* transcription by binding to the *PAL-boxes* in its promoter, we looked for any changes in metabolites derived from the phenylpropanoid biosynthetic pathway. We measured lower levels of key lignin biosynthesis intermediates in *A. rabiei*–infected chickpea compared to mock-infected plants, while the flavonoid and amino acid biosynthetic pathways remained unaffected. Additionally, the wild type and ArPEC25 RNAi transformants infected chickpea plants also showed the expected modulation of lignin biosynthesis intermediates, suggesting a direct role of ArPEC25 in the lignin biosynthetic pathway. We conclude that the biosynthesis of major lignin subunits is directly hampered during pathogenesis. The chemical composition and integrity of the lignin polymer largely depend upon the type and ratio of its constituent subunits. Lignin in conifers is predominantly composed of G subunits with a small fraction of H subunits, whereas woody dicots use mainly G and S subunits with an S/G ratio of 2 (Wang et al., 2014). It would be interesting to determine the relative biosynthesis rate of each type of lignin subunit and whether the ratios between subunit types change.

ArPEC25 harbors two cysteine residues (C-35 and C-88). ArPEC25 also homodimerizes, as revealed by Y2H assay (Supplemental Figure S6B). The cysteine residues in effectors are known to facilitate various functions during host invasion. First, they form intermolecular disulfide bonds for homodimerization. Second, cysteine residues also assist in the interactions between the effector and its host targets, as was reported for the SsSSVP1 effector of *S. sclerotiorum* (Lyu et al., 2016). Third, these inter- and intra-molecular disulfide bonds prevent the degradation of effectors in the harsh chemical environment of the host (Liu et al., 2012). It will be interesting to explore the precise function of the two cysteine residues in ArPEC25. Although ArPEC25 contains the conserved PEXEL motif that was initially characterized in *Plasmodium* spp. for its role in effector secretion and translocation from the parasite to host erythrocytes (Boddey et al., 2009, 2010, 2016), we failed to detect a role for this motif in effector secretion in our system (Figure 2B). Many fungal genomes, including that of *A. rabiei*, encode an array of effectors with this conserved PEXEL motif (Luisa Hiller et al., 2004; Choi et al., 2010; Verma et al., 2016). Whether the PEXEL motif has some unexplored function such as host entry is therefore a matter of conjecture.

*CaPAL1* transcript levels rose early during infection in the susceptible chickpea varieties Pusa362 and WR315, before returning to basal levels in later stages of infection. By contrast, *CaPAL1* transcript levels remained low during the early stages of infection of the resistant chickpea varieties ILC3279 and ILC6482, followed by a marked upregulation in the later stage of infection. Plant transcription factors such as KNOX, LIM, and MYB regulate *PAL* isogenes in different plants and may jointly regulate the tissue- and stress-specific expression patterns of these genes. LIM transcription factors are also known to form complexes with other transcription factors such as AT-HOOK MOTIF CONTAINING NUCLEAR LOCALIZED (AHL) members (Zhao et al., 2013). Surprisingly, in chickpea, *CaAHL18* is a putative candidate gene for resistance against AB within the mapping interval of the QTL *qABR4*.*1* (Kumar et al., 2018). Hence, the increased expression of Ca*PAL1* in AB-resistant chickpea varieties in the later stage of infection may reflect the association of CaβLIM1a with other such transcription factors. The activity of this transcription activator complex would be severely impeded by the higher levels of ArPEC25 resulting from the rapid growth of *A. rabiei* on the susceptible chickpea varieties Pusa362 and WR315.

In conclusion, we propose a mechanism whereby CaβLIM1a plays a role in host immunity by fortifying the physical barrier of the cell wall via lignin deposition in normal chickpea plants. The necrotrophic pathogen *A. rabiei* facilitates infection through its secreted virulence factor ArPEC25, which translocates to the host cell nucleus and modulates the DNA binding and transactivation function of CaβLIM1a via direct physical interaction (Figure 9). The impaired activity of the transcription factor results in reduced production of lignin subunits and a weakened cell wall to support successful penetration and virulence.

**Figure 9.**
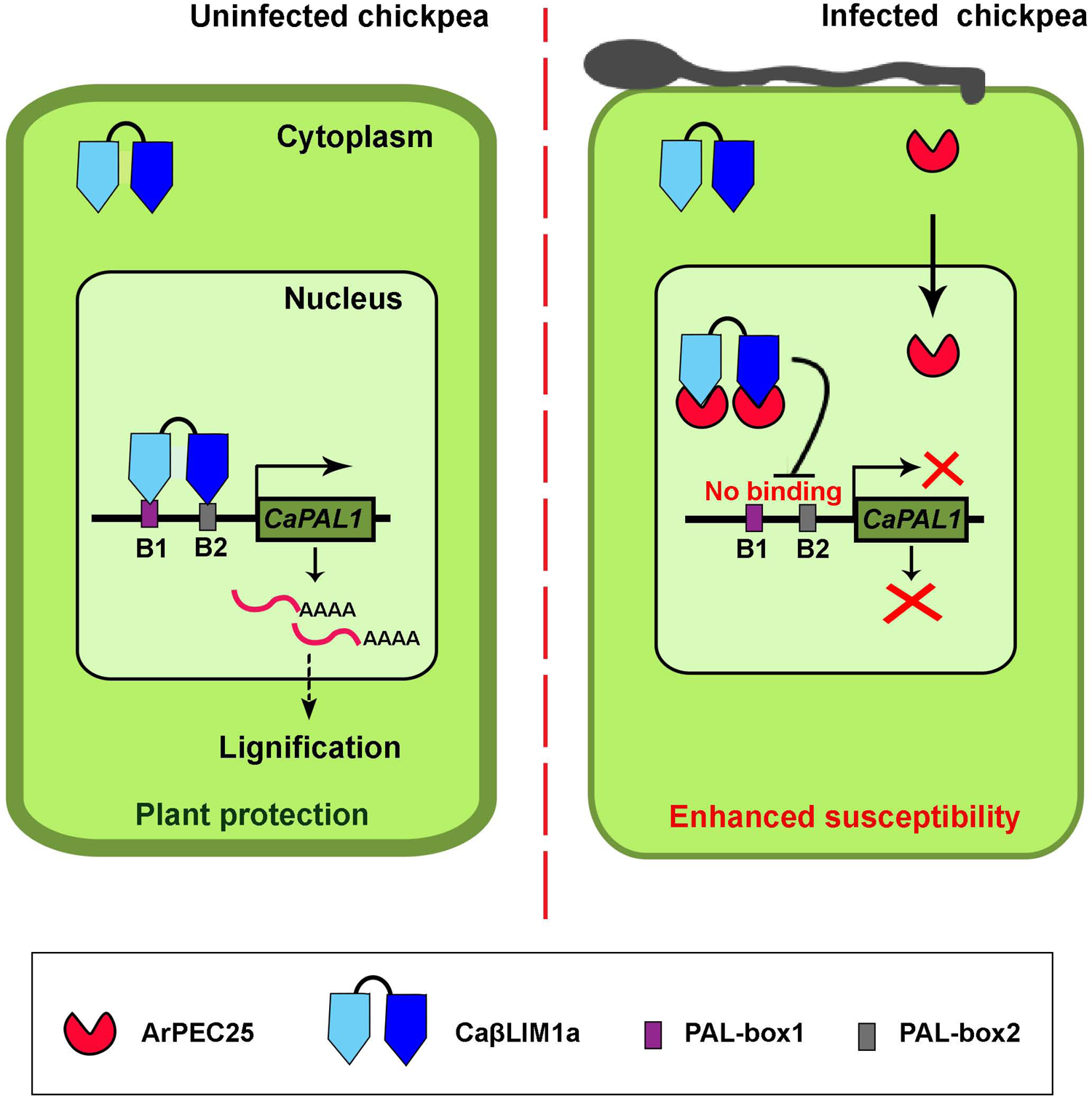
Proposed working model of the role of ArPEC25 in chickpea susceptibility. Pathogen secreted effector ArPEC25 interacts with CaβLIM1a and inhibit its binding to promoter sequence. In normal chickpea plants, the CaβLIM1a transcription factor binds directly to the promoter sequence of *CaPAL*1 gene resulting in the activation of defense related phenylalanine ammonia-lyase pathway which leads to the synthesis of lignin components. Pathogen secreted effector ArPEC25 translocate to host cell nucleus and interacts with a LIM domain containing TF, CaβLIM1a and negatively regulate the expression of *CaPAL*1 gene by inhibiting the binding of TF on *PAL-box*1 and *PAL-box*2 of *CaPAL*1 promoter. Intervention in TF binding leads to negative modulation in the expression of chickpea *CaPAL*1 gene and ultimately down regulation in the synthesis of lignin and hence compromised host.

## Methods

### Culture conditions, DNA isolation

The wild type virulent *A. rabiei* isolate ArD2 (Indian Type Culture Collection No. 4638) was obtained from the Division of Plant Pathology, Indian Agricultural Research Institute (New Delhi, India). *A. rabiei* was routinely maintained on potato dextrose agar (PDA; Difco Laboratories, pH 5.2 to 5.5) at 22°C for 15-20 days. To harvest fungal mycelia, conidia were allowed to grow in potato dextrose broth (PDB; Difco Laboratories, pH 5.2 to 5.5) at 22°C, 120 rpm for 5-7 days. Total genomic DNA from mycelial balls was isolated using a Quick-DNA™ Fungal/ Bacterial Miniprep Kit (Zymo Research, USA) as per the manufacturer’s instructions.

### RNA isolation and gene expression analysis

The total RNA from fungal mycelia, and Chickpea Pusa 362 plant tissues infected with *A. rabiei* conidia, was isolated using TRIzol reagent (Invitrogen, USA). The cDNA was synthesized using High-Capacity cDNA Reverse Transcription Kit, (Thermo Fisher Scientific, USA). Targeted gene expression was determined by real-time PCR with ABI7900 (Applied Biosystem, USA) using gene specific oligonucleotide (Supplemental table S3) and Brilliant III Ultra-Fast SYBR Green QPCR Master Mix (Agilent Technologies, USA). Relative gene expressions were calculated using 2^−ΔΔCt^ method, and the values were derived from three independent biological replicates, each having three technical replicates. *Caβ-tubulin* (LOC101495306) and elongation factor 1-alpha (*ArEF1α*; ST47_g4052) was used as an internal control for chickpea and *A. rabiei*, respectively.

## PEG mediated fungal genome editing and complementation in *A. rabiei*

To generate *ArPEC*25 knockout construct (Δ*arpec*25), 623 bp *5’* flanking regions of *ArPEC*25 was PCR amplified from *A. rabiei* genomic DNA using gene specific primer combination (Supplemental table S3). Amplified PCR product was cloned at *Xho*I and *Hind*III restriction site in pGKO2-hph vector, using In-Fusion cloning (Clontech, Japan), to generate pGKO2-5’*ArPEC*25-*hph* construct. Similarly, 1024 bp *3’* flanking sequence of *ArPEC*25 was PCR amplified using gene specific primer combination (Supplemental table S3) and cloned at *Bam*HI and *Eco*RI site in pGKO2-5’*ArPEC*25-*hph* construct using In-Fusion cloning strategy to generate the knockout construct pGKO2-5’*ArPEC*25-*hph*-3’*ArPEC*25. The cloned *ArPEC*25 gene replacement construct of ∼ 3.5 kb, including *5’* and *3’* flanking sequences and gene sequence of hygromycin (*hph*), was amplified using primer combination 5’-ArPEC25-F and 3’-ArPEC25-R, and transformed into *A. rabiei* protoplasts, as described earlier (Sinha et al., 2021).

The *ArPEC*25 complementation construct (Δ*arpec*25/*ArPEC*25), was prepared by amplifying full length *ArPEC*25 along with its native promoter and terminator, using specific primers combinations (Supplemental table S3). The amplified fragment was cloned at *Eco*RI and *Xba*I site in pBIF2-mEos2 vector (having kanamycin and G-418 selections for bacteria and fungus, respectively). Similarly, deletion mutant carrying a transgene encoding a version of ArPEC25 with a nuclear export signal (*Δarpec*25/*ArPEC25*-*NES*) was prepared using specific primer combinations (Supplemental table S3), and cloned at *Eco*RI and *Xba*I site in pBIF2-mEos2 vector using In-Fusion cloning. Complementation constructs were independently transformed into Δ*arpec*25 knock-out mutant, through *Agrobacterium tumefaciens* mediated transformation (ATMT), as described previously (Nizam et al., 2010). Knockout and complementation constructs were confirmed using Southern blot assay as described earlier (Sinha et al., 2021).

### Mini-dome bioassay

The mini-dome technique described earlier (Cho et al., 2004), was used to measure virulence of wild type *A. rabiei*, and its derived mutant Δ*arpec*25, and complementation transformants Δ*arpec*25/*ArPEC*25, and Δ*arpec*25/*ArPEC*25-NES. The chickpea plants were grown under controlled conditions at D/N temperature: 24^0^C/18^0^C; D/N light duration: 14/10; Light intensity: 250 µE/m2/s for day; Relative Humidity: 100%. Two weeks old chickpea seedlings were spray inoculated with fungal conidia (2 × 10^6^ spores/ml), collected from 20-days-old fungus. Inoculated seedlings were immediately covered with a transparent plastic cup to provide a uniform high level of relative humidity, required for infection to occur. The mini-domes were removed at 48 hours post inoculation (hpi). Disease severity was rated 7 days post inoculation (dpi).

For in-vitro oxidative stress treatment, 250 µM Menadione (Sigma-Aldrich, USA), was added to one-week old broth culture of *A. rabiei* (inoculated with 100 µl of 1 × 10^4^ conidia/ml spore suspension). Mycelial balls were harvested using three layers of Mira cloth (EMD, Millipore Corp, Germany) at 0.25 h, 0.5 h, 1 h, 3 h, and 6 h post treatment. Ethanol treated samples were used as control. For gene expression during in-planta infection, *A*. *rabiei* conidial suspensions (2 × 10^6^ conidia/ml) were spray inoculated on two-weeks old chickpea seedlings (Pusa 362). Infected stems and leaves portions were harvested at 12, 24, 72, and 144 hpi.

### Yeast Two-Hybrid Assay

The interaction between chickpea TFs CaβLIM1a, CaβLIM1b, CaδLIM2, CaWLIM1a, CaWLIM1b, CaWLIM2, and an *A. rabiei* effector ArPEC25 in yeast system was investigated using split-ubiquitin based DUAL hunter system (Dualsystems Biotech, Switzerland). Genes were cloned into bait (pGDHB1) and prey (pGPR3-N) vectors at *Not*I and *Asc*I site. The cloned plasmids along with their controls were co-transformed into yeast strain NMY51, using EZ-Yeast transformation kit (MP biomedicals, USA). Transformed yeast cells were grown on synthetic dextrose (SD, -Leu, -Trp) medium for 3 days at 30°C. Yeast clones were spotted on SD (-Leu, -Trp) medium as selection for positive clones and SD (-Leu, -Trp, -His, -Ade) medium supplemented with 15 mM 3-amino-1,2,4-triazole (3-AT) as selection for protein-protein interaction and incubated for 3 d at 30°C. The interaction between CaβLIM1a-Cub and NubI served as a positive control, while the co-expression of CaβLIM1a-Cub and NubG was considered as negative control. Primers used in the cloning are enlisted in supplemental data set (Supplemental table S3).

### Yeast one-hybrid assays

Binding of CaβLIM1a to its *cis* regulatory element on the promoter sequence of *CaPAL*1 (LOC101507594) was confirmed by Y1H assay. The native promoter and its deletion constructs were cloned into the pAbAi vector. Yeast strain Y1H gold was transformed with these constructs, and positive clones were selected on synthetic dextrose (SD) medium without Ura (-Ura) supplemented with 150 ng ml^-1^ AbA (Aureobasidin A). *CaβLIM1a* was amplified and cloned into the pGADT7 vector. The cloned plasmids, along with their controls were transformed into Y1H strain containing the promoter sequence of *CaPAL*1. Binding of CaβLIM1a on *CaPAL*1 promoter was confirmed by spotting of yeast clones on selection plate SD-Leu-Ura, supplemented with 150 ng ml^-1^ AbA. Empty vector served as negative control. Primers used in the cloning are enlisted in supplemental data set (Supplemental table S3).

### Dual-luciferase reporter assay

The 737 bp promoter sequence of *CaPAL*1 was amplified from chickpea genomic DNA, cloned in Entry vector and mobilized into destination vector which contain NLS-RFP p635NLS-RFP using Gateway LR Clonase II enzyme mix to generate *ProCaPAL*1*-FLuc* reporter construct. The reporters were co-expressed with different effector constructs such as CaβLIM1a-MYC, CaWLIM1a-MYC and ArPEC25-Flag into *N. benthamiana* leaves and empty vector (EV) served as control. Infiltrated leaf discs were collected 48 hpi, grinded to fine powder in liquid nitrogen and mixed with 1x passive lysis buffer provided in the Dual-Luciferase® Reporter Assay System (Promega, USA). Firefly luciferase and REN activity were measured following manufacture’s instruction (Promega, USA) using the POLARstar Omega multimode plate reader (BMG Labtech, Germany). The luciferase activity was calculated by normalizing the REN expression level. Immunoblotting was used to confirm the expression of effector constructs. Primers used in the cloning are enlisted in supplemental data set (Supplemental table S3).

### Transactivation assay in yeast

The open reading frame (ORF) of *CaβLIM1a* was cloned using gene specific primers (Supplemental table S3) into the pGBKT7g vector from the Entry clone using Gateway LR Clonase II enzyme mix. The construct pGBKT7gCaβLIM1a was transformed into the Y2H Gold yeast strain following EZ-Yeast transformation kit (MP biomedicals, USA). The pGBKT7:CaWRKY50 (Kumar et al., 2016) was used as positive control. Empty pGBKT7 vector served as negative control in transactivation assay. The assay was screened on selective medium (-His, -Ade).

### Subcellular localization

For subcellular localization study, proteins were transiently expressed into *N. benthamiana* leaves. The coding sequence of *ArPEC*25 without signal peptide (ArPEC25ΔSP) and *CaβLIM*1*a* were mobilized into pGWB405-35S-GFP from the respective Entry clone using Gateway LR Clonase II enzyme mix. NLS -RFP vector was used as nuclear marker. For co-localization of CaβLIM1a and ArPEC25, the ArPEC25ΔSP was cloned into pCAMBIA1301-mCherry vector. The constructs were transiently co-expressed in *N. benthamiana* leaves. The fluorescence signal was detected 48 hpi using TCS SP8 confocal laser scanning microscope (Leica Microsystems, Germany). Primers used in cloning are enlisted in supplemental data set (Supplemental table S3).

### Recombinant Protein Preparation

To construct the plasmid for recombinant protein production, the ArPEC25ΔSP was fused with Strep II by primer-based amplification, and cloned in pET28a (+) vector to create pET-PEC25-Strep II-6xHis. The recombinant proteins were expressed in BL21 cells with 1 mM isopropyl β-D-thiogalactoside (IPTG) for 6 h at 28°C, and affinity purified with Strep-Tactin® Sepharose® beads (iba, Germany) following manufacturer’s instructions. For the purification of MBP-CaβLIM1a or MBP alone, transformed bacterial cells were induced with 1 mM IPTG for 4 h at 28°C and expressed proteins were affinity purified with Amylose Resin (Biolabs, New England). Primers used in cloning are enlisted in supplemental data set (Supplemental table S3).

### Electrophoretic Mobility Shift Assay (EMSA)

Gene encoding *CaβLIM1a* was amplified and cloned into pMALC2x vector to generate MBP-CaβLIM1a recombinant protein. The resulting construct was then transformed into BL21, and the recombinant MBP-CaβLIM1a was purified as mentioned above. Purified MBP alone served as control. Three different DNA probe containing *PAL-boxes* was synthesized, dimerized and labeled with [γ-^32^P] ATP as per the protocol of Gel Shift Assay System (Promega, USA). For EMSA, the purified MBP/MBP-CaβLIM1a was mixed with the radio labeled probe and 5x binding buffer (from respective Gel Shift Assay System kit) and incubated for 20 min at room temperature. The reaction was electrophoresed on 6% polyacrylamide gel in 0.5x TBE (pH 8.3), and transferred to a phosphor screen and imaged in Typhoon® (GE Healthcare, USA).

### Yeast secretion trap (YST) assay

The pYST1 vector containing ArPEC25 with and without SP was transformed in *suc2* yeast mutant strain. Preparation of chemo-competent yeast cells and transformation by the high-efficiency lithium acetate (LiAc) method was performed according to Clontech yeast transformation user manual. The transformants were selected on synthetic dextrose -Leu (SD/-L) medium for 3 to 5 days. The transformed colonies from the primary selection plates were then re-suspended in liquid SD/-L and grown to stationary phase overnight at 30°C and 250 rpm. Aliquots of these cultures were diluted in distilled water to a density of OD_600_ 1. Spotting for secretion analysis was performed on SD/-L medium containing 2% sucrose and antimycin A at 2 μg/mL.

### Bimolecular Fluorescence Complementation Assay

The ORF of *CaβLIM*1*a* was cloned in frame with the N-terminal half of Venus (CaβLIM1a-NVenus) and ArPEC25ΔSP was translationally fused to C-terminal of Venus (ArPEC25ΔSP-CVenus) in pDOE vector (Gookin and Assmann 2014). NLS-RFP was used as nuclear marker. The constructs were transformed into *Agrobacterium* GV3101 and co-infiltrated in *N. benthamiana* leaf. YFP signal was detected using TCS SP8 confocal laser scanning microscope (Leica Microsystems, Germany) 48 hpi. Presence of mTurquoise2 signal indicates successful transient expression of respective genes in the leaf. Primers used in cloning are enlisted in supplemental data set (Supplemental table S3).

### Protein extraction and western blot

Total protein was extracted from 0.2 g of infiltrated fresh *N. benthamiana* leaves by grinding to fine powder in liquid nitrogen. Powder was then resuspended in 1x PLB provided in the Dual-Luciferase® Reporter Assay System (Promega, USA). Extract was vortexed and then centrifuged at 16,000 g for 1 min at 4°C. Supernatant was collected for protein gel blot analysis.

For total protein extraction from yeast, 2 mL culture of yeast cells were harvested through centrifugation and resuspended in 2 M Lithium acetate (LiAc). Cells were further harvested by centrifugation and then resuspended in 0.4 M NaOH for 5 min on ice. Harvested cells were finally resuspended in protein loading dye boiled for 5 min and supernatant collected for immunoblotting.

For total protein extraction from fungal mycelia, fungal conidia were grown to mycelial balls in PDB,and treated with 250 µM Menadione (Sigma-Aldrich, USA) for 3 h and harvested by passing through three layers of sterile Mira cloth (EMD, Millipore Corp, Germany). Fungal tissue was grounded to fine powder in liquid nitrogen and homogenized in Tris-glycine buffer pH 8.3 (3 g Trizma and 14.4 g Glycine, Sigma-Aldrich, USA in 1 L MQ). Lysate was centrifuged at 16,000 g for 40 min at 4°C. The supernatant was collected for protein gel blot analysis with specific antibody. For total secreted protein extraction from culture filtrate (CF), PDB grown 7 days old fungal mycelia was treated with 250 µM Menadione (Sigma-Aldrich, USA) for 3 h. Axenic CF were separated from fungal mycelia by passing it through three layers of Mira cloth as mentioned above. Further, CF was sequentially filtered with 0.45 and 0.22-micron filter disc (Durapore PVDF, Sigma-Aldrich, USA). Filtered CF was then concentrated using 3 kDa Amicon® Ultra-15 Centrifugal Filter Units (Merck, USA). The concentrate was then snap freezed and stored in −80°C for future use.

Proteins were separated by SDS-PAGE and transferred to polyvinylidene fluoride (PVDF) Transfer Membrane, (MDI, India), followed by overnight blocking at 4°C in 2% Non-Fat Powdered Milk, (Bio Basic, Canada INC.) in 1x Phosphate Buffer Saline (PBS) supplemented with 0.1% TWEEN20 (Sigma-Aldrich, USA). For detection, membrane was incubated with primary antibody Myc (Catalog CPA9004), HA (Catalog CPA 9002), Flag (Catalog CPA9001) from Cohesion Biosciences, Anti-Old Yellow Enzyme 6 (Anti-Oye6) and anti-His, with 1:2000 dilution for 1 h at room temperature. Gt anti-Rb IgG (H+L) Secondary antibody, HRP conjugate (Catalog 65-6120; Invitrogen, USA) with 1:5000 dilutions was used for 1 h at room temperature.

Detection was performed by Clarity^TM^ Western ECL substrate (BIO-RAD, USA).

### Pull Down and protein identification through mass spectrometry

For the immuno purification of fungal secreted ArPEC25, *A. rabiei* was transformed with ArPEC25-FLAG construct using ATMT. Conidia from the stable transformants were allowed to grow in PDB. The axenically grown mycelial balls were separated, and the CF was concentrated as mentioned above. The concentrated CF was used for immuno purification of ArPEC25-FLAG using anti-Flag antibody coated Dynabeads™ Protein G, (ThermoFisher, USA), and purified following manufacturer’s instructions. The purified protein was precipitated and dissolved in G-buffer. 25 µL samples were taken and reduced with 5 mM TCEP and further alkylated with 50 mM iodoacetamide and then digested with trypsin (1:50, trypsin/lysate ratio) for 16 h at 37°C. Digests were cleaned using a C18 silica cartridge and dried using a speed vac. The dried pellet was resuspended in buffer A (5% acetonitrile, 0.1% formic acid).

Processed samples were identified by mass spectrometric analysis using EASY-nLC 1000 system (ThermoFisher Scientific, USA) coupled to QExactive mass spectrometer (ThermoFisher Scientific, USA) equipped with nano electrospray ion source. About 1.2 µg of the peptide mixture was resolved using a 15 cm PicoFrit column (360 µm outer diameter, 75 µm inner diameter, 10 µm tip) filled with 1.9 µm of C18-resin (Dr Maeisch, Germany). The peptides were loaded with buffer A and eluted with a 0–40% gradient of buffer B (95% acetonitrile, 0.1% formic acid) at a flow rate of 300 nl/min for 45 min. MS data was acquired using a data-dependent top10 method dynamically choosing the most abundant precursor ions from the survey scan. For data processing, three reaction were processed and 3 RAW files generated were analyzed with Proteome Discoverer against the ArPEC25 sequence. For Sequence search, the precursor and fragment mass tolerances were set at 10 ppm and 0.5 Da, respectively. The protease used to generate peptides, i.e. enzyme specificity was set for trypsin/P (cleavage at the C terminus of “K/R: unless followed by “P”) along with maximum missed cleavages value of two. Carbamidomethyl on cysteine as fixed modification and oxidation of methionine and N-terminal acetylation were considered as variable modifications for database search. Both peptide spectrum match and protein false discovery rate were set to 0.01 FDR.

### Gel Filtration Assay

Purified proteins were subjected to gel filtration analysis using Superdex 200 Increase 10/300 GL column (GE Healthcare Bio-Sciences AB) with a flow rate of 0.2 mL/min and injection volume of 2 mL and fraction size of 0.5 mL. Phosphate buffer saline (Sigma-Aldrich, USA) was used for the assay. The eluted fractions were analyzed through western blot with α-His Ab following SDS-PAGE. For the gel filtration assay of mixture of MBP-CaβLIM1a and ArPEC25-6xHis, proteins were purified and incubated in equal amount for 30 min at room temperature and were subjected to gel filtration assay as described above.

### Quantitative measurement and statistical analysis

Lesion size and diameter of AB infection were measured using ImageJ/Fiji software. FRET efficiency was calculated in Leica software. Statistical significance of means or difference among groups were obtained from the average of three biological replicates, having at least three technical replicates in each group. To calculate statistical significance Student’s t-test and ANOVA followed by Tukey test between the multiple groups, were performed using GraphPad Prism 6. The *p* < 0.05 was accepted as significant.

### *In silico* bioinformatics analysis

All the gene and protein sequences in this study were obtained from NCBI (http://www.ncbi.nlm.nih.gov). Multiple sequence alignment was performed by PROMALS3D using default parameters and then visualized by Jalview.. Molecular evolution and phylogenetic studies of ArPEC25 was performed through MEGA X software using neighbor-joining method, having bootstrap value of 1000. To determine the putative TFs binding site on the promoter sequence of *CaPAL*1, PlantCARE online tools (Lescot et al., 2002) was used. To find out the theoretical pI of the protein Expasy compute pI/Mw tool was used.

### RNAi

For the downregulation of *ArPEC*25, RNAi strategy was used. For RNAi, pSilent I (Nakayashiki et al., 2005) vector was modified, and cloning of *ArPEC*25 ORF was performed in sense and antisense directions separated by cutinase intron. *A*. *rabiei* was transformed with pSilent I: *ArPEC*25 through ATMT as described earlier. Putative *ArPEC*25 RNAi transformants were screened for the downregulation of the gene through real-time PCR, as described earlier. Further, specific transformant was used for metabolite studies.

### Metabolite analysis

Chickpea (Pusa 362) plants were infected with conidia of *A*. *rabiei* or *ArPEC*25 RNAi as mentioned above. The samples were harvested at 24 hpi and 72 hpi, and stored in liquid nitrogen for downstream processing. The samples were completely desiccated into lyophilizer. Next, lyophilized samples were grounded to fine powder in tissue lyser, and ∼100 mg dry weight sample was used for metabolite extraction. For the total extraction of metabolite, 80% methanol (HPLC grade) was added to the fine powder followed by heating at 65°C for 15 min. The samples were incubated at 28°C for overnight for complete extraction of different metabolites followed by centrifugation at 10,000 rpm for 5 min, to remove cell debris. The supernatant was collected and vaporized in speed vac at room temperature. The remained pellet was re-dissolved in 1 ml 80% methanol and filtered through 0.45-micron filter disc (Durapore PVDF, Sigma-Aldrich, USA). UHPLC was performed using 50 µl of this filtered sample and the data was analyzed.

## Author contribution

PKV conceptualized the work and SKS, SV, KS, AS, RS VS, KK performed the experiments, AP, SKS, PKV analyzed the data, SKS, and PKV wrote the manuscript and funding managed by PKV.

## Acknowledgment

Authors acknowledge the financial support by the Department of Biotechnology, Government of India through research grant for the Challenge Program on Chickpea Functional Genomics Project (Sanction No. BT/AGR/CG-Phase II/01/2014), and core grant from National Institute of Plant Genome Research (NIPGR), New Delhi, India. SKS acknowledges Department of Biotechnology, Government of India for a fellowship.

## Supplementary figure legends

**Supplemental Figure 1.** Knockout of *ArPEC*25 (Supports Figure 1). **(A)** Schematic diagram of homologous recombination based *ArPEC*25 deletion strategy. *HPH* stands for hygromycin phosphotransferase gene. **(B)** Southern hybridization to confirm *ArPEC25* knockout. Band on the blot confirms the successful gene deletion and single integration of *hph* at replacement site. 1; wild type, 2; *Δarpec25*, 3; *Δarpec*25/*ArPEC*25 and 4; *Δarpec*25/*ArPEC25*-*NES*. **(C)** Disease symptoms on AB susceptible chickpea plants after 7days post inoculation. Pusa362 plant showed necrotic lesions on stem and leaves after infection with *A. rabiei* (WT) and indicted fungal transformants. **(D)** Relative expression of *ArPEC25* in WT and indicated fungal transformants. Real-time PCR result of *ArPEC25* in WT, knock-out (*Δarpec25*), complementation strain (*Δarpec*25/*ArPEC*25) and mutant carrying a transgene encoding a version of ArPEC25 with a nuclear export signal (*Δarpec*25/*ArPEC25*-*NES*). The error bar represents the mean ± SD value. Transcript abundance of *ArEF1a* was used as internal control. Statistically significant differences were calculated by unpaired one tailed t-test; * P ≤ 0.0388. **(E)** Hyphal germination of WT and *Δarpec*25 as seen under confocal microscopy. No noticeable deference in germination pattern between *Δarpec*25 and WT was observed. Bar, 10 µm.

**Supplemental Figure 2.** Evolutionary relationships of ArPEC25 with other homologous proteins. The tree was constructed using sequences from: *Ascochyta rabiei*, *Epicoccum nigrum* (OSS44957.1), *Curvularia kusanoi* (KAF3003731.1), *Didymella heteroderae* (KAF3039597.1), *Didymella keratinophila* (KAF3042860.1), *Macroventuria anomocheta* (XP 033561316.1), *Didymella exigua* (XP 033448406.1), *Ascochyta lentis* (KAF9694412.1), *Lizonia empirigonia* (KAF1350522.1), *Polyplosphaeria fusca* (KAF2728976.1), *Dothidotthia symphoricarpi* (XP 033523869.1), *Ophiobolus disseminans* (KAF2829292.1), *Stagnospora sp.* (OAK93469.1), *Parastagnospora nodorum* (QRC97135.1), *Pyrenochaeta sp.* (OAL45347.1), *Cucurbitaria berberidis* (XP 040783831.1), *Pyrenophora acminiperda* (RMZ72407.1), *Pyrenophora teres* (CAA9981104.1), *Pyrenophora tritici-repentis* (XP 001939465.1), *Bipolaris oryzae* (XP 007692785.1), *Bipolaris sorokiniana* ND90Pr (XP 007694363.1), *Bipolaris sorokiniana* (KAF5847504.1), *Stemphylium lycopersici* (R845.1), *Alternaria atra* (CAG5163760.1), *Alternaria burnsii* (XP 038781177.1), *Alterneria alternata* (XP 018383193.1), *Alterneria arborescens* (RYO72432.1), *Alternaria arborescens* (XP 028506601.1), *Plenodomus tracheiphilus* (KAF2845459.1). The multiple sequence alignment was performed by PROMALS3D software and phylogenetic tree was constructed suing MEGAX software. The bootstrap value derived from 1000 iterations.

**Supplemental Figure 3.** Multiple sequence alignment of ArPEC25 protein with its homologs from various organisms. The alignment was generated with the multiple sequence and structure alignment program PROMALS3D using default parameters and then visualized by Jalview. The positions of the conserved PEXEL motif and arginine-rich region are highlighted with rectangular boxes. The two conserved cysteine residues present at the N- and C-terminal are indicated by asterisk. A stretch of ten residues, RECPVPKPGG at the C-terminal shown in rectangular box was also highly conserved. Shaded boxes below the alignment indicate the degree of conservation.

**Supplemental Figure 4.** Screening of native and PEXEL mutant transformants (Support Figure 2B). **(A)** Schematics representation of native and PEXEL mutant used for transformation. The alanine (A) in red represents substitution of sequence in the motif. The effector domain represents the remaining sequences of ArPEC25. C-terminal Flag and YFP tag is also shown. (B) Confocal microscopy used for screening of fungal transformants. ArPEC25 with RTLND sequence and mutant variants as shown in **(A)** were expressed in *A. rabiei*. YFP signal was detected under confocal microscopy. Bar, 5 µm. (C) PEXEL mutant transformants screened by immuno blot using anti-Flag antibody. Molecular weights of the proteins are shown on the right with black arrow. Ponceau stain (lower panel) indicates the presence of protein in all lanes. “+” and “-” indicate presence and absence, respectively.

**Supplemental Figure 5.** Screening of positive over expression (OE) transformants and protein immuno-precipitation (Support Figure 3C, D and E) **(A)** Immuno blot of OE transformants tissue lysate. The positive OE transformant was screened using anti-Flag antibody. Ponceau stain indicate presence of protein in all lane. WT tissue lysate served as a control. **(B) and (C)** Protein immuno purification from processed CF used in (Figure 3C). **(B)** SDS PAGE of processed CF of OE transformants. **(C)** Same CF was also used for the immuno precipitation of ArPEC25 using Flag antibody for mass spectrometric analysis. Purified ArPEC25 is shown with black arrow head. **(D)** MS based analysis for cleavage of RTLND sequence in secreted ArPEC25. The CF of OE transformant was collected and processed for purification of ArPEC25 using Flag antibody. Purified protein was subjected to LC-MS/MS for detection of cleavage of RTLND at L residue. Intact PRTLNDSAD detected in the spectra represents un cleaved motif.

**Supplemental Figure 6.** Localization study of ArPEC25 (Support Figure 2A) and its homodimerization in Y2H assay. **(A)** Localization study of GFP tagged ArPEC25 in *N. benthamiana*. Indicated ArPEC25-GFP-GUS and ArPEC25ΔSPAAAAAAA-GFP-GUS were infiltrated with nuclear marker NLS-RFP in leaf cells. Confocal image indicates that ArPEC25-GFP-GUS localizes to nucleus (upper panel), whereas, mutant ArPEC25ΔSPAAAAAAA-GFP-GUS missed nucleus (lower panel). Bar, 50 µm. **(B)** ArPEC25 form homodimer in Y2H assay. Yeast cells containing indicated plasmids were grown in indicated selective medium. Growth of cells on interaction selective SD (-LWAH) medium suggests homodimerization of ArPEC25. The Large T and Δp53 and pAI-Alg5 and pDL2-Alg5 are positive controls and autoactivation, respectively. SD (-LW), synthetic dextrose (-Leu, Trp) medium; SD (-LWAH), synthetic dextrose (-Trp, Leu, His, Ade) medium. **(C)** Confocal microscopy of fungal transformant treated with and without H_2_O_2_. Indicated translational fusion constructs were expressed in wild type *A. rabiei* and microscopy was performed for H_2_O_2_ treated and untreated samples. Bar, 5 µm. ‘+’ and ‘–’ represent presence and absence.

**Supplemental Figure 7.** Interaction of ArPEC25 with Ca2LIMs (Support Figure 5A and 5G). **(A)** Full image of Figure 5A showing interaction of ArPEC25 with all six CaLIM transcription factors. **(B)** Confocal image showing expression of donor and acceptor proteins used in FRET experiment. The donor CaβLIM1a-GFP and the acceptor ArPEC25-mCherry were transiently co-expressed in *N. benthamiana* leaves. Images were taken 48 hpi using confocal microscopy for pre-bleach and post bleach samples (upper panel). Lower panel represents the enlarged image of the section outlined by white dashed line in (upper panel). White arrow head indicate the nucleus considered for FRET. Bar, 20 µm.

**Supplemental Figure 8.** Schematics of vector used in the study. **(A)** Schematic diagram of constructs used for yeast secretion trap assay. ArPEC25 with and without SP are cloned independently in translational frame of *Suc*2 gene under the regulation of P*ADH*1. **(B)** Schematic diagram of constructs used for FRET assay. For FRET donor, CaβLIM1a is translationally fused to GFP. In FRET acceptor, effector ArPEC25 is translationally fused to mCherry. **(C)** Schematic diagram for domain deletion in CaβLIM1a (Support Figure 6C). The LIM1 and LIM2 domain of CaβLIM1a has been sequentially deleted. **(D)** Schematic diagram of the promoter of *CaPAL1* (Support Figure 6D). The 737 bp promoter of *CaPAL1* shows the presence of *PAL-box1* and *PAL-box2* and their sequential deletion.

**Supplemental Figure 9.** Transactivation of CaLIMs, and expression of *CaPAL* genes. **(A)** Transactivation of CaLIMs in the yeast system. CaWLIM1b, CaβLIM1a, CaβLIM1b, CaWLIM2, CaWLIM1a and CaδLIM2, respectively were tested for activator property. Indicated plasmids were expressed in yeast. Growth of cells on indicated selective medium suggests transactivation. SD/(-W), synthetic dextrose (-Trp) medium, SD/(-H), synthetic dextrose (-His) and SD/(-W-H-A), synthetic dextrose (-Trp, His, Ade). **(B)** Western blot of constructs used in Figure 5C. Cell lysate of *N. benthamiana* expressing EV, *CaPAL*1-CaβLIM1a and *CaPAL*1-CaWLIM1a used for the luciferase assay in Figure 5C were taken for immuno blot. Expressions of proteins were detected with anti-Myc antibody. Red asterisk represents band of CaβLIM1a. Ponceau stain (lower panel) indicates the presence of protein in all lanes. **(C)** Repressor assay in yeast system. All six CaLIMs used for transactivation assay in S. Figure 9A were tested for the repressor activity. The activation domain VP16 was used for the experiment. Empty vector, pBRIDGE was used as negative control. **(D)** Expression level of *CaPAL*1, *CaPAL*2, *CaPAL*3 and *CaPAL*4 in *A. rabiei* infected chickpea cultivar Pusa362 at different time point (0, 3, 6, 12, 24, 72, 144 and 216 h). The relative expression values were determined by real-time PCR and the error bar represents mean ± SD of three technical replicates from an independent experiment. Statistically significant differences were calculated by Two-way Anova, Dunnett’s multiple comparisons test.

**Supplemental Figure 10.** Promoter binding assay for CaLIM TFs. **(A)** Promoter binding of all six CaLIM TFs in Y1H assay. Yeast strain containing indicated plasmids were grown on selection medium. Growth of cells in selection medium supplemented with indicated concentration of AbA suggests DNA protein interaction. Black triangle represents increasing dilution of yeast cells. SD/(-U), synthetic dextrose (-Ura), SD/(-L), synthetic dextrose (-Leu), SD/(-L^AbA150^), synthetic dextrose (-Leu supplemented with150 ng/ml aureobasidin A). **(B)** Immuno blot of yeast transformants used in S. Figure 10A. Y1H strain transformed with indicated constructs mentioned in S. Figure 10A were used for immuno blot. Anti-HA antibody was used for the detection of band. **(C)** Immuno blot in yeast (Support Figure 6D). Yeast strain expressing CaβLIM1a with full length *CaPAL*1 promoter or *PAL-box* deletion variants were used for immuno blot. Anti-HA antibody was used for detection of band. Ponceau stain (lower panel) indicates the presence of protein in all lanes. “+” and “-” indicate presence and absence, respectively. **(D)** Immuno blot of truncated CaβLIM1a (Support Figure 6C). Yeast strain expressing full length CaβLIM1a and different LIM deletion variant used in Figure 6C were taken for immuno blot. Anti-HA antibody was used for the detection of band. Ponceau stain (lower panel) indicates the presence of protein in all lanes. **(E)** Purification of recombinant protein used in EMSA Figure 6F and 7A. Left panel shows the purified MBP and MBP-CaβLIM1a while the right panel shows the purified ArPEC25-6xHis.

**Supplemental Figure 11.** Knockdown of *ArPEC*25. **(A)** Schematics of the construct used for gene knockdown in wild type *A. rabiei*. The sense and antisense strand of full length *ArPEC*25was cloned under the control of promoter *TrpC* and was used for *Agrobacterium* mediated transformation of wild type *A. rabie*. **(B)** Transcript level of *ArPEC*25 is shown in different RNAi transformants (#22 and #34) and wild type treated with 250 µM menadione for 3 h. Gene transcript abundance was determined by real-time PCR and samples were normalized by fungal *ArβTUB* mRNA. The data shown for fold change represents the mean ± SD value of three technical replicates from three independent experiments (n=3). Statistically significant differences were calculated by unpaired two tailed t-test; **** P ≤ 0.0001 and *** P ≤ 0.0007. **(C)** Hyphal germination of wild type *A. rabiei* and RNAi transformant (#22) as seen under confocal microscopy. No noticeable deference in the germination pattern of knock-down transformant compared to WT was observed Bar, 10 µm.

**Supplemental Figure 12.** Schematics of pathway of various metabolites in their corresponding branches of phenylpropanoid biosynthetic pathway and their differential accumulation revealed by heat map. Metabolites marked with red showed significant differential regulation in UHPLC assay. Phenyl ammonia lyase (PAL) enzyme is shown in blue color. Heat map of the corresponding branch of the pathway shows the log2 transformed values of the concentration of metabolites in dried chickpea tissues treated with mock and *A. rabiei*. The colors of the heat map for metabolite levels represents their average fold-change values. The data used to make the heat-map is average of at least three biological replicates (detailed statistical analysis is given in Supplemental Figure S13 and S14).

**Supplemental Figure 13.** Content of key metabolites of phenylpropanoid pathway synthesizing flavonoids and anthocyanin in mock and *A. rabiei* treated chickpea. Metabolic compounds were quantified by separating methanolic extracts from dried tissues of chickpea treated with mock and wild type *A. rabiei* using UHPLC. The graph shows the ± SD of at least three independent treated chickpea plants. Statistically significant differences were calculated by unpaired student t-test.

**Supplemental Figure 14.** Modulation in the accumulation of different amino acids in chickpea treated with mock and *A. rabiei*. Accumulation data of amino acids are recorded for at least three independent biological replicates using UHPLC spectrum of methanolic extract of dried chickpea tissues treated with mock and wild type *A. rabiei* and plotted as fold change with respect to mock treated sample. Error bar represents ± SD value. Statistically significant differences were calculated by unpaired student t-test.

## Supplementary tables

Supplemental table S1

Summary of candidate *A. rabiei* effector proteins containing conserved sequence motifs.

Supplemental table S2

List of putative ArPEC25 interacting transcription factors found in chickpea library screening.

Supplemental table S3

List of primers used in the study.

